# Elucidating the kinetic and thermodynamic insight into regulation of glycolysis by lactate dehydrogenase and its impact on tricarboxylic acid cycle and oxidative phosphorylation in cancer cells

**DOI:** 10.1101/2024.06.26.600909

**Authors:** Siying Zeng, Yuqi Wang, Minfeng Ying, Chengmeng Jin, Chang Ying, Di Wang, Hao Wu, Xun Hu

## Abstract

Lactate dehydrogenase (LDH) stands at the intersection of pyruvate metabolism. While it is believed that inhibition of LDH redirects pyruvate to mitochondrial metabolism, suppressing glycolysis and boosting oxidative phosphorylation, the mechanism remains largely unexplored. We found that individual LDH A or B knockouts had minimal impact on glycolysis, tricarboxylic acid cycle (TCA cycle), or oxidative phosphorylation (OXPHOS). However, combining LDH knockout with LDH inhibitor GNE-140 significantly suppressed these processes. Inhibition of LDH led to an increase in free NADH concentration and a decrease in free NAD^+^ concentration, the reduced free NAD^+^ concentration inhibited GAPDH, disrupting the balance of glycolytic intermediates, which were linked with thermodynamic shift of the Gibbs free energy of reactions between phosphofructokinase 1 (PFK1) and phosphoglycerate mutase (PGAM) in the glycolytic pathway, favoring their reverse direction. This disrupted glycolysis led to impaired TCA cycle and mitochondrial respiration due to reduced pyruvate and glutamine carbon influx into TCA cycle. Under hypoxia, LDH inhibition had a stronger effect, inducing energy crisis, redox imbalance, and cancer cell death. Our study reveals LDH’s intricate control over glycolysis, TCA cycle, and mitochondrial respiration, highlighting the interplay of enzyme kinetics and thermodynamics in metabolic pathways, a crucial aspect for understanding metabolic regulation.

**Impact statement:** This study elucidates a biochemical mechanism by which lactate dehydrogenase influences glycolytic flux in cancer cells, revealing a kinetic– thermodynamic interplay that contributes to metabolic regulation.

## Introduction

A prominent metabolic feature of most cancer cells is Warburg effect or aerobic glycolysis, characterized by high glycolytic rates accompanied with excessive production of lactate even with ample oxygen (Liberti & Locasale, 2016; Vander Heiden, Cantley, & Thompson, 2009; Warburg, 1956). LDH catalyzes the final step of the glycolytic pathway, converting NADH (generated at the step of GAPDH) and pyruvate (generated at the step of pyruvate kinase) into lactate and NAD^+^. In cancer cells, the actual change in the Gibbs free energy (ΔG) of the LDH-catalyzed reaction ranges between -3 to -7 kJ/mol (Jin, Zhu, Wu, Wang, & Hu, 2020; X. Zhu, Jin, Pan, & Hu, 2021), favoring lactate accumulation. Consistently, lactic acidosis (high lactate concentration and acidic pH) is common in the microenvironment of most solid tumors (Dantas et al., 2016; Gatenby & Gillies, 2004; Peppicelli, Bianchini, & Calorini, 2014; Perez-Tomas & Perez-Guillen, 2020; Raghunand, Gatenby, & Gillies, 2003). Lactic acidosis plays multiple roles in tumor progression, including but not limited to immune suppression (Choi, Collins, Gout, & Wang, 2013), promotion of angiogenesis (Fukumura et al., 2001), enhancement of metastasis (Walenta et al., 2000), regulation of metabolism (Li et al., 2022). Clinically, lactic acidosis and high LDH expression are associated with poor prognosis of cancer patients (Augoff, Hryniewicz-Jankowska, & Tabola, 2015; Cui et al., 2014; McCleland et al., 2012; Petrelli et al., 2015; Walenta et al., 2000). Thus, inhibiting LDH is a potential way to treat cancer and numerous LDH inhibitors have been developed for this purpose (Boudreau et al., 2016; Le et al., 2010; Maftouh et al., 2014; Manerba et al., 2012; Rai et al., 2017; Z. Zhao, Han, Yang, Wu, & Zhan, 2015).

LDH is located at a critical point in pyruvate metabolism, where it regulates pyruvate flow to lactate and to mitochondrial respiration. Previous studies have shown that inhibition of LDH leads to suppression of glycolysis while enhancement of mitochondrial respiration in cancer cells, and it is believed that inhibition of LDH directs pyruvate towards mitochondrial metabolism. Fantin et al. demonstrated that knockdown of LDH-A using short hairpin RNAs reduced LDH activity, decreased lactate production, and stimulated mitochondrial respiration (Fantin, St-Pierre, & Leder, 2006). Le et al. reported that inhibition of LDH-A activity by siRNA knockdown and LDH inhibitor FX-11 were accompanied by enhanced mitochondrial oxygen consumption and increased mitochondrial reactive oxygen species, leading to oxidative stress (Le et al., 2010). LDH inhibition-induced oxidative stress further led to cancer cell death (Le et al., 2010). Zdralevic et al. showed that genetic disruption of both LDHA and LDHB or inhibition of LDH by GNE-140, a potent LDH inhibitor, shifted cancer cells from glycolysis to oxidative phosphorylation (Zdralevic et al., 2018). These studies suggest that LDH acts as a critical regulator balancing ATP production between glycolysis and OXPHOS: when LDH activity is high, pyruvate generated from glycolysis is mainly converted to lactate, resulting in low mitochondrial respiration; when LDH activity is low, pyruvate generated from glycolysis is directed towards mitochondrial metabolism, increasing mitochondrial respiration.

Despite the progress made in understanding the potential role of LDH as described above, there exists an unresolved critical issue: the exact biochemical mechanism behind how it affects glycolysis and mitochondrial respiration isn’t fully understood. If the effect of LDH inhibition were confined solely to its catalytic step, as suggested in previous reports, this could explain enhanced mitochondrial respiration by directing pyruvate towards mitochondrial metabolism. However, this explanation conflicts with other facts. Notably, the glycolytic rate is far higher than the mitochondrial respiration rate (Liberti & Locasale, 2016). According to the glycolytic rate and OXPHOS rate in different cancer cells (Zeng & Hu, 2023), we estimate that the ratios of glycolytic rate over oxidative phosphorylation rate are 10.53 to 47.11. Thus, mitochondria can only consume a minor fraction of pyruvate generated from glycolysis, this cannot explain the markedly reduced lactate generation rate. Moreover, the activity of LDH in cancer cells is far higher than that of hexokinase (HK), the first rate-limiting enzyme in the glycolytic pathway (Jin et al., 2020; J. Xie, Dai, & Hu, 2016; X. Zhu et al., 2021). According to calculation of our published works (Jin et al., 2020; J. Xie et al., 2016; X. Zhu et al., 2021), even if LDH activity is inhibited by 90%, the remaining LDH activity is still significantly higher than that of HK and would be sufficient to convert pyruvate generated from glycolysis to lactate. Therefore, inhibition at the LDH’s catalytic step may not account for the significant decrease in overall lactate production. We previously reported that glycolysis rate was regulated by both enzyme kinetics and chemical thermodynamics within the glycolytic pathway (Jin et al., 2020; X. Zhu et al., 2021). Based on the above reasoning, we propose the following hypothesis: upon LDH inhibition, via not yet deciphered mechanisms involving enzyme kinetics and chemical thermodynamics in the glycolytic pathway, the rate at the upstream segment of the glycolytic pathway is suppressed, leading to reduced rates of glycolysis. Moreover, with the overall rate of pyruvate production reduced and the remaining LDH activity sufficient to convert pyruvate to lactate, there would be a decrease in pyruvate flux to mitochondrial respiration, leading to a suppression of mitochondrial respiration. If this hypothesis holds true, it would provide a more complete understanding of how LDH inhibition impacts cancer cell metabolism and could provide new clues for the development of more effective therapeutic strategies. Because therapeutic inhibition of LDH is expected to act over sustained periods, we focus on the metabolic steady state re-established after LDH suppression rather than on the earliest transient responses immediately following perturbation.

## Results

### The effect of LDHA or LDHB knockout on glycolysis

The first question to address is if LDHA or LDHB knockout would significantly affect the glycolytic rate. We determined the relationship between LDH activity and glycolysis rate in HeLa-LDHAKO (LDHA knockout), HeLa-LDHBKO (LDHB knockout), and HeLa-Ctrl cells. The relative LDH activities of HeLa-LDHAKO and HeLa-LDHBKO cells were about 30% and 70% of the control, indicating LDHA and LDHB accounted for about 70% and 30% of the total LDH activity, respectively (Figure. 1A). Despite significant differences in LDH activity, the glucose consumption rate and lactate production rate were comparable between 3 groups (Figure. 1B). We repeated the above experiments using the mouse breast cancer cell 4T1 model and obtained similar results, i.e., knockout of LDHA or LDHB did not significantly affect glucose consumption and lactate production (Figure 1—figure supplement 1A & B). The reason may be that the total activity of LDH in HeLa cells is very high, which is 297-fold higher than the first rate-limiting enzyme HK activity in HeLa-Ctrl cells and 81-fold higher in HeLa-LDHAKO cells (Table supplement 1). This enzyme activity ratio is common in other tumor cells (Jin et al., 2020; J. Xie et al., 2016; X. Zhu et al., 2021).

**Figure 1.**
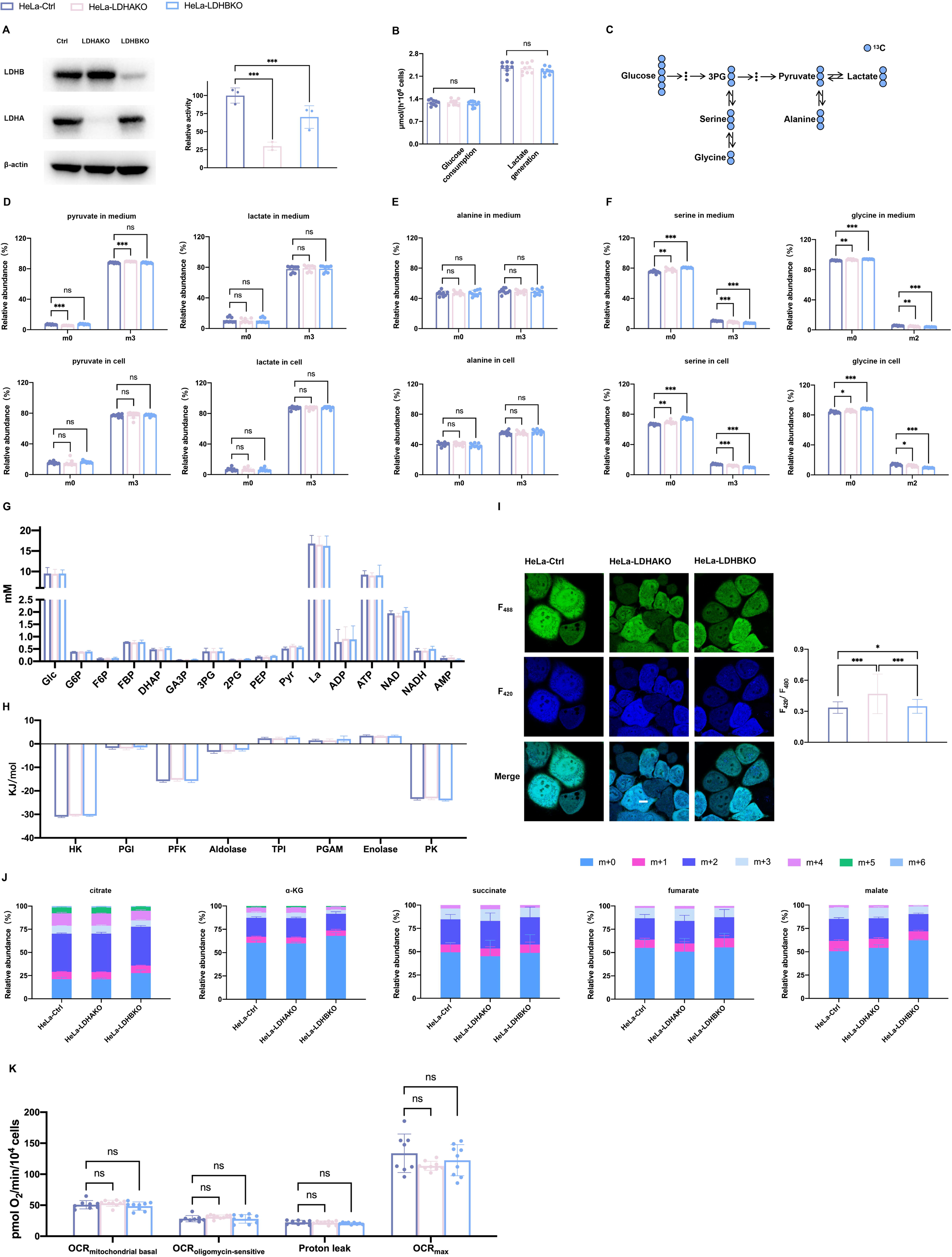
The effects of LDHA or LDHB knockout on LDH activity, glycolysis, glucose carbon into TCA cycle and OXPHOS in HeLa cells. (A) Representative Western Blot of LDHA and LDHB and relative LDH activity in the cell lysate of HeLa-Ctrl, HeLa-LDHAKO, and HeLa-LDHBKO. (B) Glucose consumption rate and lactate generation rate. Cells were cultured in complete RPMI-1640 medium in a CO_2_ incubator for 6 hours, and then the medium concentrations of glucose and lactate were determined as described in Materials and Methods. (C-F) Tracing glucose carbon to pyruvate, lactate, alanine, serine, and glycine. Cells were cultured in complete RPMI-1640 medium containing 6 mM [^13^C_6_]glc in a CO_2_ incubator for 6 hours, and then the percentages of isotopologues of pyruvate, lactate, alanine, serine, and glycine in cells and in medium were determined by LC-MS/MS as described in Materials and Methods (Table supplement 13). (G) The concentrations of glucose, lactate, and the glycolytic intermediates in cells (Table supplement 14). (H) ΔG of the reactions in the glycolytic pathways (Table supplement 15). (I) Free NADH/NAD^+^ in cells represented by ratiometric SoNar. Representative fluorescent confocal microscopic images of cells transfected with SoNar (left) and the statistics (right). Scale bar: 10μm. (J) Tracing glucose carbon to TCA cycle intermediates (citrate, α-KG, succinate, fumarate, malate). Cells were cultured in complete RPMI-1640 medium containing 6 mM [^13^C_6_]glc in a CO_2_ incubator for 6 hours, and then the percentages of isotopologues of the TCA cycle intermediates in cells were determined by LC-MS/MS as described in Materials and Methods. (K) OCR, measured as described in Materials and Methods. Data are mean ± SD from 3 independent experiments. *, *P*<0.05, **, *P*<0.01, ***, *P*<0.001.

The above results demonstrated that neither LDHAKO nor LDHBKO significantly affected glucose conversion to lactate in HeLa cells, and the results were different from previous reports (Arseneault et al., 2013; Q. Chen et al., 2022; X. Chen, Liu, Kang, Gnanaprakasam, & Wang, 2023; H. Xie et al., 2014). In addition to glycolytic rate, we sought to investigate the effect of LDHAKO or LDHBKO on the glycolytic pathway in the following four aspects:

#### 1. The glucose carbon to pyruvate and lactate

We used [^13^C_6_]glc to trace glucose conversion to pyruvate and lactate (Figure 1C). This approach could rule out interference from lactate production by other metabolic pathways, e.g., glutaminolysis (DeBerardinis et al., 2007; Perez-Escuredo et al., 2016). The results showed that the isotopologues percentages of pyruvate and lactate were comparable between 3 groups, both in culture medium and in cells (despite being statistically significant, the difference in terms of m3 isotopologues% of pyruvate in medium was very small, 1.81%) (Figure 1D). The isotope tracing results in line with the quantification of lactate confirmed that glucose conversion to pyruvate and lactate through glycolysis was not significantly affected.

#### 2. The glucose carbon to subsidiary branches of glycolysis

We traced glucose carbon to subsidiary branches of glycolysis (Figure 1C). Pyruvate can be converted to alanine through alanine aminotransferase. The isotopologues of alanine (m0 and m3) were similar between 3 groups (Figure 1E), indicating that the inhibition of LDH did not significantly affect the proportion of glucose-derived pyruvate converting to alanine. We traced [^13^C_6_]glc-derived serine, which was produced from 3PG, a glycolytic intermediate. The m0 serine species was provided by the culture medium, and the m3 serine isotopologue was generated from [^13^C_6_]glc through 3PG via serine synthesis pathway. Serine can be converted to glycine by serine hydroxymethyl transferase. There were statistically significant yet moderately decreases of m3 serine% and m2 glycine% in HeLa-LDHAKO and HeLa-LDHBKO cells (Figure 1F), indicating LDHAKO or LDHBKO moderately reduced 3PG to serine synthesis pathway.

#### 3. The concentration of glycolytic intermediates and the thermodynamic state of the pathway

Then, we measured the concentrations of glycolytic intermediates in the three groups of cells. The concentrations of these intermediates did not change significantly (Figure 1G), neither the ΔGs of the reactions in the glycolytic pathway (Figure 1H), indicating that LDHAKO or LDHBKO did not significantly alter the thermodynamic state in the glycolytic pathway.

#### 4. The redox state of cytosolic free NADH/NAD^+^

Because LDHAKO or LDHBKO markedly reduced the total LDH activity, in theory, it would lead to an increase in its substrate concentration. As pyruvate concentrations were comparable between 3 groups of cells, NADH concentration may increase significantly in HeLa-LDHAKO and HeLa-LDHBKO cells. NAD^+^ and NADH in cells exist in 2 forms, bound and free (Xiao, Wang, Handy, & Loscalzo, 2018; X. H. Zhu, Lu, Lee, Ugurbil, & Chen, 2015). Although LDHAKO or LDHBKO exerted no significant effect on total concentrations of NAD^+^ and NADH (Figure 1G), we speculated that there would be an effect on free NAD^+^ and NADH. We measured the intracellular free NADH/NAD^+^, represented by the ratiometric fluorescent SoNar probe (Y. Zhao et al., 2015; Y. Zhao et al., 2016), which showed an increase of free NADH/NAD^+^ (Figure 1I). The increase of free NADH/NAD^+^ was more significant in HeLa-LDHAKO cells, as expected. The results suggested that the decrease in total LDH concentration was counterbalanced by a concomitant increase in the concentration of its substrate, free NADH, thereby maintaining the reaction velocity.

Similarly, LDHA or LDHB KO did not significantly affect the concentrations of intermediates in the glycolytic pathway in 4T1 cells (Figure 1—figure supplement 1C, table supplement 2), nor did it significantly change ΔGs in the glycolytic pathway (Figure 1 — figure supplement 1D, table supplement 3), but it significantly affected free NADH/NAD^+^ (Figure 1—figure supplement 1E).

Together, LDHAKO or LDHBKO did not significantly affect glycolysis, including glucose conversion to lactate, the concentrations of glycolytic intermediates, and the thermodynamic state of glycolytic pathway (the delta Gs of the reactions in the glycolytic pathway). However, LDHAKO or LDHBKO significantly increased the cytosolic free NADH/NAD^+^ and significantly decreased glycolytic intermediates shuttling to serine synthesis pathway.

### The effect of LHDAKO or LDHBKO on contribution of glucose carbon to TCA cycle and on mitochondrial respiration

Because LDHAKO or LDHBKO did not significantly affect the rate of glycolysis, nor it altered the proportion of pyruvate to lactate, we speculated they would not significantly affect glucose-derived pyruvate entering into TCA cycle. To confirm it, we used [^13^C_6_]glc to trace glucose carbon entering into TCA cycle. The labeling percentage of the TCA cycle intermediates (citrate, α-KG, succinate, fumarate, malate) by glucose carbon, in general, were comparable between HeLa-Ctrl, HeLa-LDHAKO, and HeLa-LDHBKO (Figure 1J). Further, we measured mitochondrial respiration of these cells, which showed no significant difference (Figure 1K). Similarly, another set of cells (4T1-Ctrl, 4T1-LDHAKO, and 4T1-LDHBKO) also showed similar level of mitochondrial respiration (Figure 1—figure supplement 2). Collectively, genetic disruption of LDHA or LDHB did not significantly affect glucose-derived pyruvate entering into TCA cycle, neither significantly affect mitochondrial respiration, although statistically significant but minimal changes were observed in a few specific parameters (e.g., m3-pyruvate% in medium), as opposed to the current point of view (Fantin et al., 2006; Le et al., 2010; Oshima et al., 2020; H. Xie et al., 2014).

### The effect of LDHBKO and GNE-140 on glycolysis - the kinetic and thermodynamic insight into the regulation of glycolysis by LDH

The above results demonstrated that LDHAKO or LDHBKO were insufficient to achieve a significant inhibition of the overall rate of glycolysis, suggesting that further LDH inhibition was required. We then used GNE-140, a potent pharmacological inhibitor of LDH (Boudreau et al., 2016). The Ki of GNE-140 on LDHA in HeLa-LDHBKO cell lysate and LDHB in HeLa-LDHAKO cell lysate were 0.88 and 9.58 μM, respectively (Figure 2A & B), indicating that the inhibitory potency of GNE-140 toward LDHA was far greater than that toward LDHB. Consistently, GNE-140 inhibited glycolysis in HeLa-LDHBKO cells significantly stronger than HeLa-LDHAKO and HeLa-Ctrl (Figure 2C). There was no significant difference between HeLa-LDHAKO and HeLa-Ctrl in response to GNE-140 (Figure 2C), and this was supported by the inhibition curve of GNE-140 on LDH activity, which was generated based on the Ki values of GNE-140 on LDHA and LDHB (Figure 2D). We checked the effects of GNE-140 on other glycolytic enzymes and found no significant inhibition nor activation of GNE-140 on other glycolytic enzymes, this excluded the inhibitory effect of GNE-140 on glycolysis via acting on other glycolytic enzymes (Figure 2—figure supplement 1).

**Figure 2.**
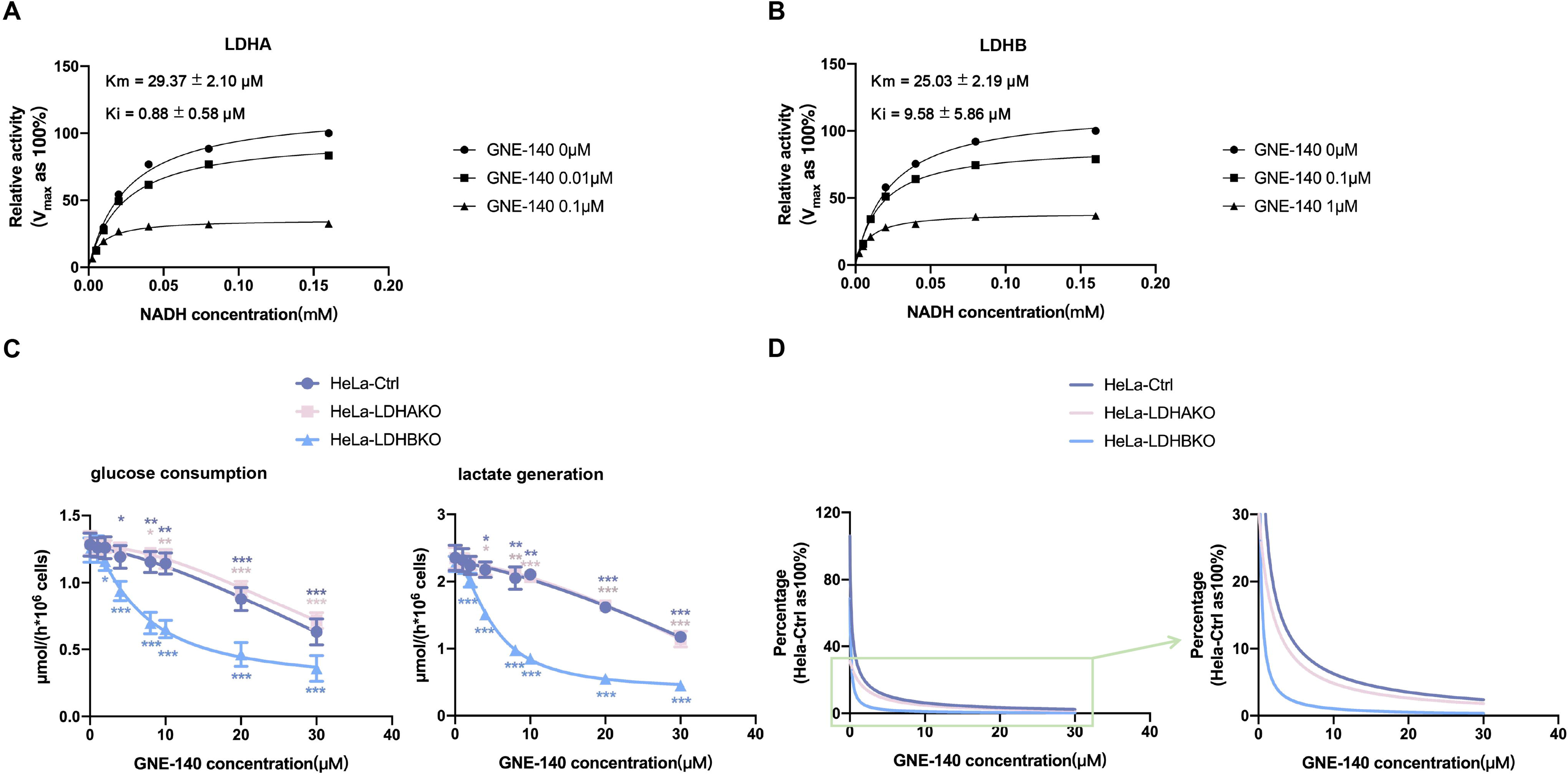
The effect of GNE-140 on LDH activity and glycolysis in HeLa cells. (A & B) Ki of GNE-140 toward LDHA and LDHB. The Ki values were determined as described in Materials and Methods. (C) The effect of GNE-140 on cellular glucose consumption and lactate generation. Cells were cultured in complete RPMI-1640 medium in a CO_2_ incubator for 6 hours, and then the medium concentrations of glucose and lactate were determined as described in Materials and Methods. (D) The curves of LDH activity versus GNE-140 concentration, which is generated based on Ki values. Data are mean ± SD from 3 independent experiments. *, *P*<0.05, **, *P*<0.01, ***, *P*<0.001.

Unlike the HeLa cell model, the Ki values of LDHA and LDHB to GNE-140 in 4T1 cells were comparable to each other (3.37 and 3.51 μM, respectively) (Figure 2 — figure supplement 2A & B). Consistently, the inhibitory effect of GNE-140 on cellular glucose consumption and lactate production was solely associated with the total activity of LDH (Figure 2—figure supplement 2C). This lack of differential effect of GNE-140 on LDHA and LDHB resulted in a distinct inhibitory pattern induced by GNE-140 in the 4T1 model compared to the HeLa model (Figure 2C).

The results in Figure 1I demonstrated that LDHAKO or LDHBKO could affect free NADH/NAD^+^ in cells to varying degrees, suggesting that free NADH/NAD^+^ was inversely proportional to the total activity of LDH. Since LDHA was far more sensitive to GNE-140 than LDHB (Figure 2A-D), treatment of HeLa-LDHBKO cells (which only express LDHA) with GNE-140 could better reflect the relationship between free NADH/NAD^+^ and total LDH activity. The results showed that there was indeed a positive relationship between GNE-140 concentrations and free NADH/NAD^+^ (Figure 3A). In line with the data in Figure 2C, LDH-inhibition-induced rate decrease of glucose consumption and lactate generation were correlated with free NADH/NAD^+^ (Figure 3B).

**Figure 3.**
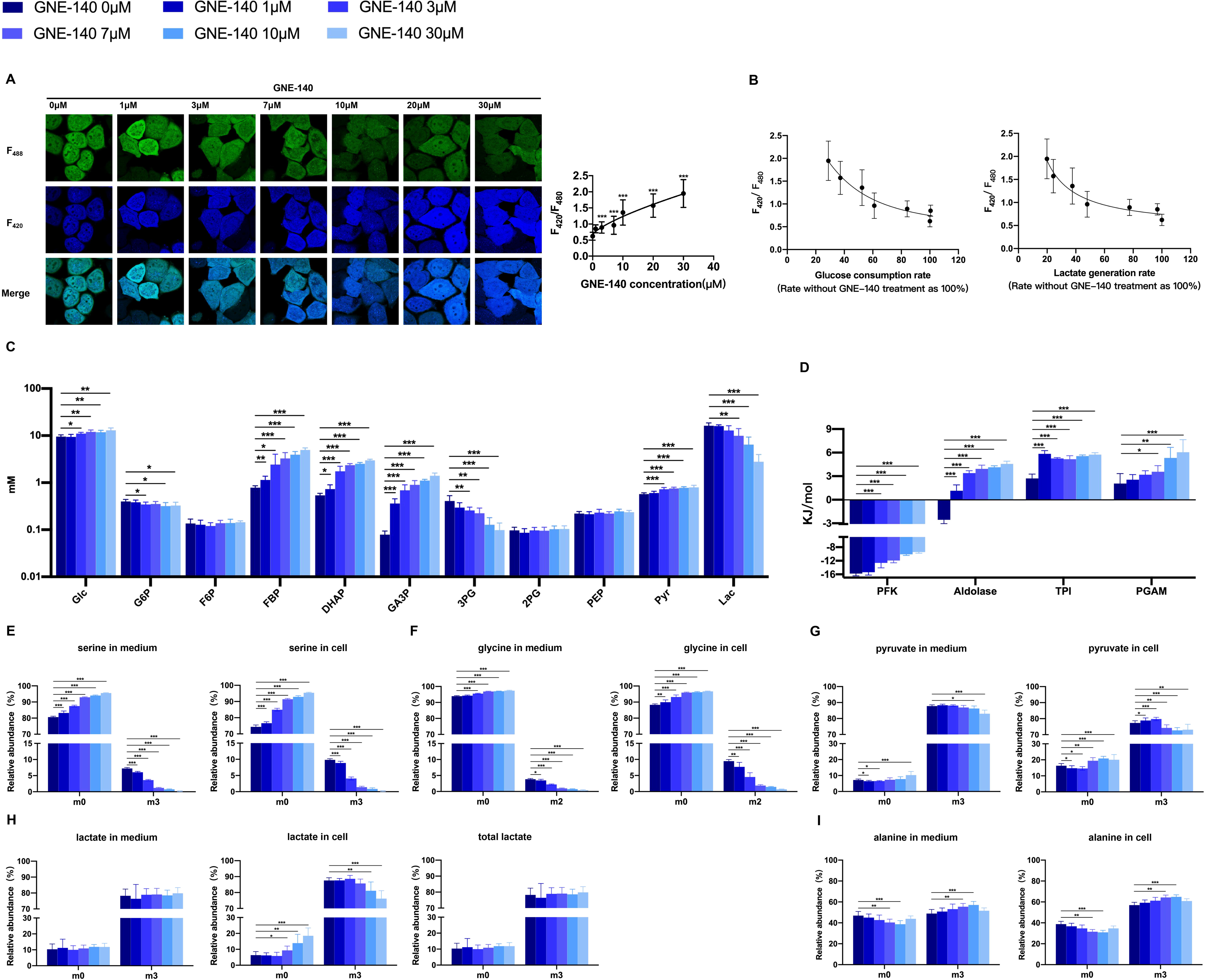
The effect of GNE-140 on the glycolytic pathway in HeLa-LDHBKO cells. (A) The effect of GNE-140 on the free NADH/NAD^+^. Representative fluorescent confocal microscopic images of cells transfected with SoNar (left) and the response of free NADH/NAD^+^ to GNE-140 (right). Scale bar: 10μm. (B) The relationship between the free NADH/NAD^+^ and glycolytic rate. (C) The effect of GNE-140 on the concentrations of glucose, lactate, and the glycolytic intermediates in cells (Table supplement 9). (D) The effect of GNE-140 on the ΔG of reactions in the glycolytic pathways (Table supplement 16). (E-I) Tracing glucose carbon to serine, glycine, pyruvate, lactate, and alanine. Cells were cultured in complete RPMI-1640 medium containing 6 mM [^13^C_6_]glc with or without GNE-140 in a CO_2_ incubator for 6 hours, and then the percentages of isotopologues were determined by LC-MS/MS as described in Materials and Methods (Table supplement 17). Data are mean ± SD from 3 independent experiments. *, *P*<0.05, **, *P*<0.01, ***, *P*<0.001.

We hypothesized that the pool of free NAD^+^ and free NADH in cells was a constant, and hence an increase in free NADH/NAD^+^ implied that the concentration of free NADH increased while the concentration of free NAD^+^ decreased. The decrease of free NAD^+^ concentration might inhibit the activity of GAPDH in the upstream of the glycolytic pathway. If so, we would observe that, with the increase in free NADH/NAD^+^ or with the increase in GNE-140 concentration, the concentration of intermediates including GA3P, DHAP, and FBP upstream of GAPDH would increase accordingly. Indeed, the concentrations of GA3P, DHAP, and FBP increased with the increase of GNE-140 concentration (Figure 3C). The results supported that LDH inhibition-induced rate decrease of glycolysis was at least partly mediated through the regulation of free NADH/NAD^+^, which modulated the activity of GAPDH.

The significant changes in the concentrations of glycolytic intermediates (FBP, DHAP, GA3P, and 3PG) were accompanied with a significant change of the ΔGs of the reactions catalyzed by PFK1, aldolase, TPI, and PGAM (Figure 3D), i.e., the PFK1-catalyzed reaction became less exergonic, aldolase-catalyzed reaction switched from exergonic to endergonic, the reactions catalyzed by TPI and PGAM were more endergonic. In essence, the changes of these ΔGs indicated more favorable for the reverse direction of the reactions, thus contributing to rate decrease of glycolysis. Supporting this, Park et al (Park et al., 2019) reported that the changes of the free energy in the glycolytic pathway, especially the reactions at the near-equilibrium state shifting to more exergonic direction, could significantly increase the rate of glycolysis, or conversely, the reactions at the near-equilibrium state shifting to more endogonic direction would significantly decrease the rate of glycolysis.

The above results suggested that the inhibition of glycolysis by GNE-140 converged in the segment between PFK1 and PGAM in the glycolytic pathway. To further confirm it, we used [^13^C_6_]glc to trace serine synthesis through 3PG in the glycolytic pathway. If the decreased rate of glycolysis was mainly at the upstream of the pathway, the rate of 3PG production would decrease, so would the rate from 3PG to serine and glycine. The results showed that intracellular m0 serine% increases, while m3 serine% decreases with the increase of GNE-140 concentration (Figure 3E). The change of m0 glycine% and m2 glycine% were the same as those of serine (Figure 3F).

Conversely, if LDH inhibition leads to glycolysis suppression at the LDH’s catalytic step, we would see a positive correlation between m3 pyruvate% and GNE-140 concentration, whereas m3 lactate% would decrease with the increasing concentration of GNE-140, as previous reported (H. Xie et al., 2014; Yeung et al., 2019). In medium, m3-pyruvate% did not change significantly until GNE-140 concentration increased to 30 μM, where m3-lactate % did not change significantly (Figure 3G). In cells, m3-pyruvate% and m3-lactate% both decreased with the increase of GNE-140 concentration, but the magnitude of decrease was more pronounced in m3-lactate% (Figure 3H). However, as medium lactate constitutes the major part, the m3-lactate% in total lactate pool (medium lactate + cellular lactate) remained not significantly changed with or without GNE-140 treatment (Figure 3H). Glucose-derived pyruvate can also convert to alanine. Intracellular and extracellular m3 alanine% remained nearly constant (Figure 3I). Taken together, the results indicated that LDH inhibition did not markedly change the proportion of pyruvate flux to lactate nor alanine.

We also traced serine, glycine, pyruvate, lactate, and alanine in HeLa-Ctrl and HeLa-LDHAKO cells by using [^13^C_6_]glc and obtained similar results (Figure 3 — figure supplement 1, table supplement 4).

Altogether, the above results pointed to that GNE-140-mediated glycolytic suppression converged at the upstream segment of the glycolytic pathway between PFK1 and PGAM.

Under hypoxia, since oxygen is deprived, cells rely more on glycolysis for ATP production. Glycolytic rate increased significantly as compared with that under normoxia (Figure 4A), and, consistently, GNE-140 was more effective with respect to inhibition of glycolysis under hypoxia than under normoxia (Figure 4A). If GNE-140-mediated glycolytic suppression was through the free NADH/NAD^+^, perturbation of GAPDH, and the Gibbs free energy of the reactions by the PFK1, aldolase, TPI, and PGAM, we should see a more significant change of these parameters. Unfortunately, we were not able to measure free NADH/NAD^+^ under hypoxia due to the technical limitations, because we could not maintain the hypoxia level under the assay condition for observing free NADH/NAD^+^. We measured the concentrations of glycolytic intermediates. There was a large change of concentrations in FBP, DHAP and GA3P (Figure 4B), more pronounced than that under normoxia condition (Figure 3C), indicating that GAPDH was inhibited. With the increase of GNE-140 concentrations, the ΔG of the reaction catalyzed by PFK1 changed from -12.68 kJ/mol to -2.56 kJ/mole, indicating that the very exergonic reaction was approaching to a near equilibrium state (Figure 4C). The ΔGs of the reactions catalyzed by aldolase, TPI, and PGAM were changing to the more endergonic state, in response to increasing concentration of GNE-140 (Figure 4C).

**Figure 4.**
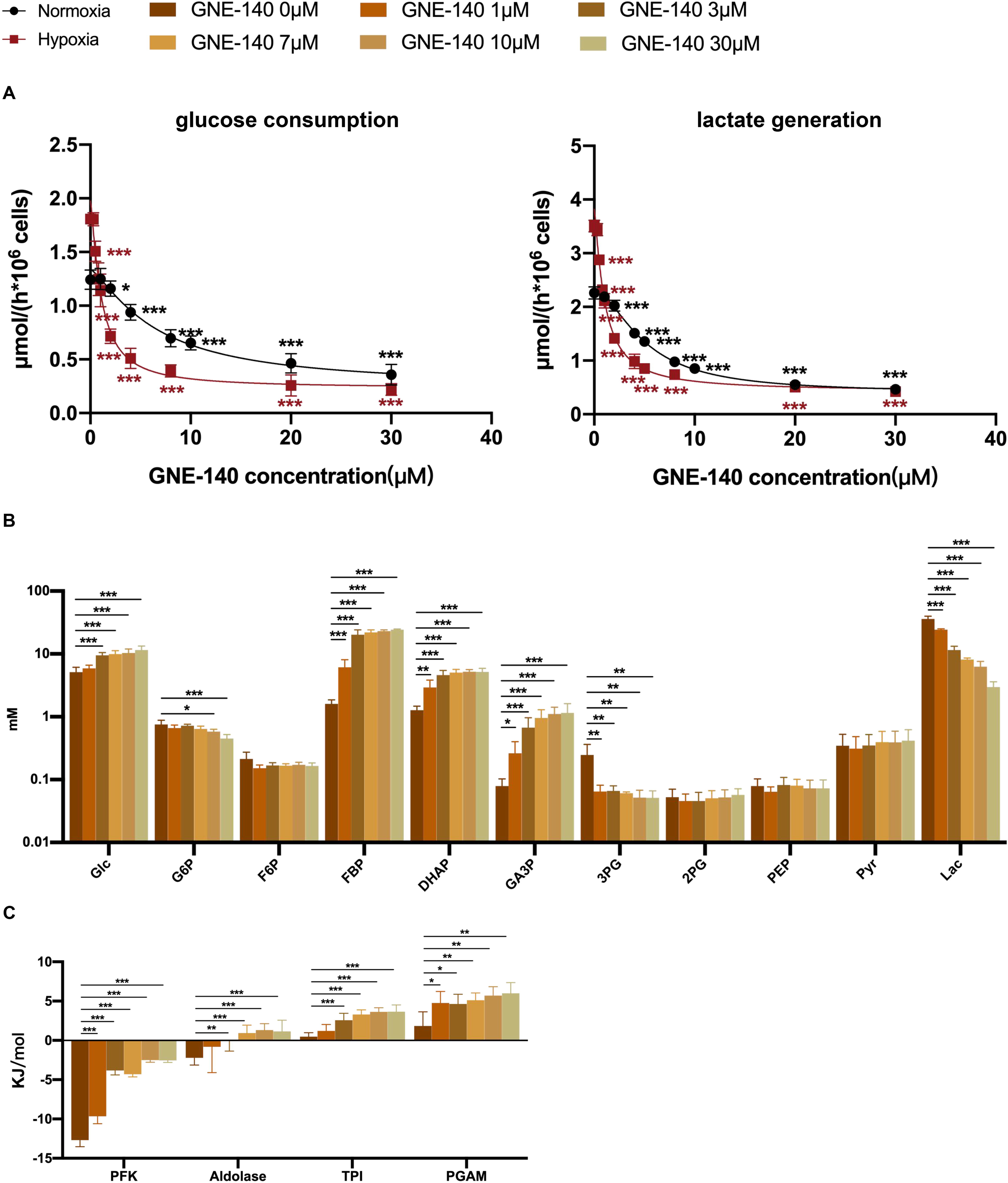
The effect of GNE-140 on the glycolytic pathway in HeLa-LDHBKO cells under hypoxia. (A) The glucose consumption rate and lactate generation rate in cells under normoxia and hypoxia (1% oxygen) and the response to GNE-140. (B) The effect of GNE-140 on the concentrations of glucose, lactate, and the glycolytic intermediates in cells under hypoxia (Table supplement 10). (C) The effect of GNE-140 on the ΔG of the reactions in the glycolytic pathways under hypoxia (Table supplement 18). Data are mean ± SD from 3 independent experiments. *, *P*<0.05, **, *P*<0.01, ***, *P*<0.001.

4T1-LDHAKO treated with GNE-140 under normoxia and hypoxia yield similar results as Hela-LDHBKO, including the increase of free NADH/NAD^+^ (Figure 3—figure supplement 2A), the inverse correlation between free NADH/NAD^+^ and glucose consumption/lactate generation rate (Figure 3—figure supplement 2B), the change of concentrations of FBP, DHAP, and GA3P (Figure 3—figure supplement 2C, Figure 4—figure supplement 1B, table supplement 5 & 6), the changes of the ΔGs of the reactions catalyzed by PFK1, aldolase, TPI, and PGAM (Figure 3—figure supplement 2D, Figure 4—figure supplement 1C, table supplement 7 & 8), and a comparison of changes in glycolysis rate with or without GNE-140 under normoxia and hypoxia (Figure 4—figure supplement 1A).

Collectively, the results offer mechanistic insights into how LDH regulates glycolysis, involving dynamic interactions between kinetics and thermodynamics. Inhibition of LDH triggers an increase in free NADH concentration while decreasing free NAD^+^ concentration, consequently perturbing GAPDH. This perturbation of GAPDH further increases concentrations of GA3P, DHAP, FBP, and decreases the concentration of 3PG, which are linked to the thermodynamic state of reactions catalyzed by PFK1, aldolase, TPI, and PGAM, favoring the reverse direction of these reactions. These sequential events contribute to the inhibition of glycolysis.

### The effect of LDHB KO and GNE-140 on the contribution of glucose carbon to TCA cycle and on OXPHOS

In Figure 2 and corresponding text, we demonstrated the sensitivity of glycolysis in HeLa-LDHBKO cells to GNE-140. We observed that GNE-140 predominantly inhibited upper segment of the glycolytic pathway, leading to a decrease in overall pyruvate production. Additionally, the proportions of glucose-derived pyruvate converting to lactate remained similar regardless of GNE-140 treatment (Figure 3H), indicating comparable proportion of the glucose-derived pyruvate reaching the mitochondria regardless of GNE-140 treatment. These results suggested that inhibition of LDH would decrease rather than increase glucose-derived pyruvate to mitochondrial metabolism. To further investigate this, we used [^13^C_6_]glucose to trace its carbon into TCA cycle. Surprisingly, the total ^13^C labeling of the TCA cycle intermediates increased with higher concentrations of GNE-140 (Figure 5A-E). When [^13^C_6_]glucose -derived pyruvate is converted to acetyl-CoA, it condenses with OAA to form citrate, generating the m2 isotopologue of citrate (Figure 5F). Citrate is the first intermediate in the TCA cycle and can be used to measure the efficiency of glucose carbon into TCA cycle. m2 citrate % decreased with the increase of the concentration of GNE-140 (Figure 5G). As m2-citrate arose from the initial incorporation of the labeled pyruvate into the TCA cycle, a decrease in the percentage of m2-citrate indicated a decrease in the direct incorporation of glucose-derived acetyl-CoA into the TCA cycle. On the other hand, other citrate isotopologues% (m1, m3, m4, m5, and m6) increased (Figure 5G). These isotopologues represented the progressive incorporation of labeled carbons from glucose-derived acetyl-CoA into the intermediates of the TCA cycle through repeated cycle, hence an increase in the percentages of these isotopologues indicated an increase in the cycling of TCA cycle intermediates or a decrease in the flux of intermediates leaving the TCA cycle. However, the percentage of m2 isotopologues of other TCA cycle intermediates (α-KG, succinate, fumarate, and malate) increased with the increased concentration of GNE-140 (Figure 5H-K). Thus, the pattern of glucose carbon labeling of citrate was different from that of other TCA cycle intermediates.

**Figure 5.**
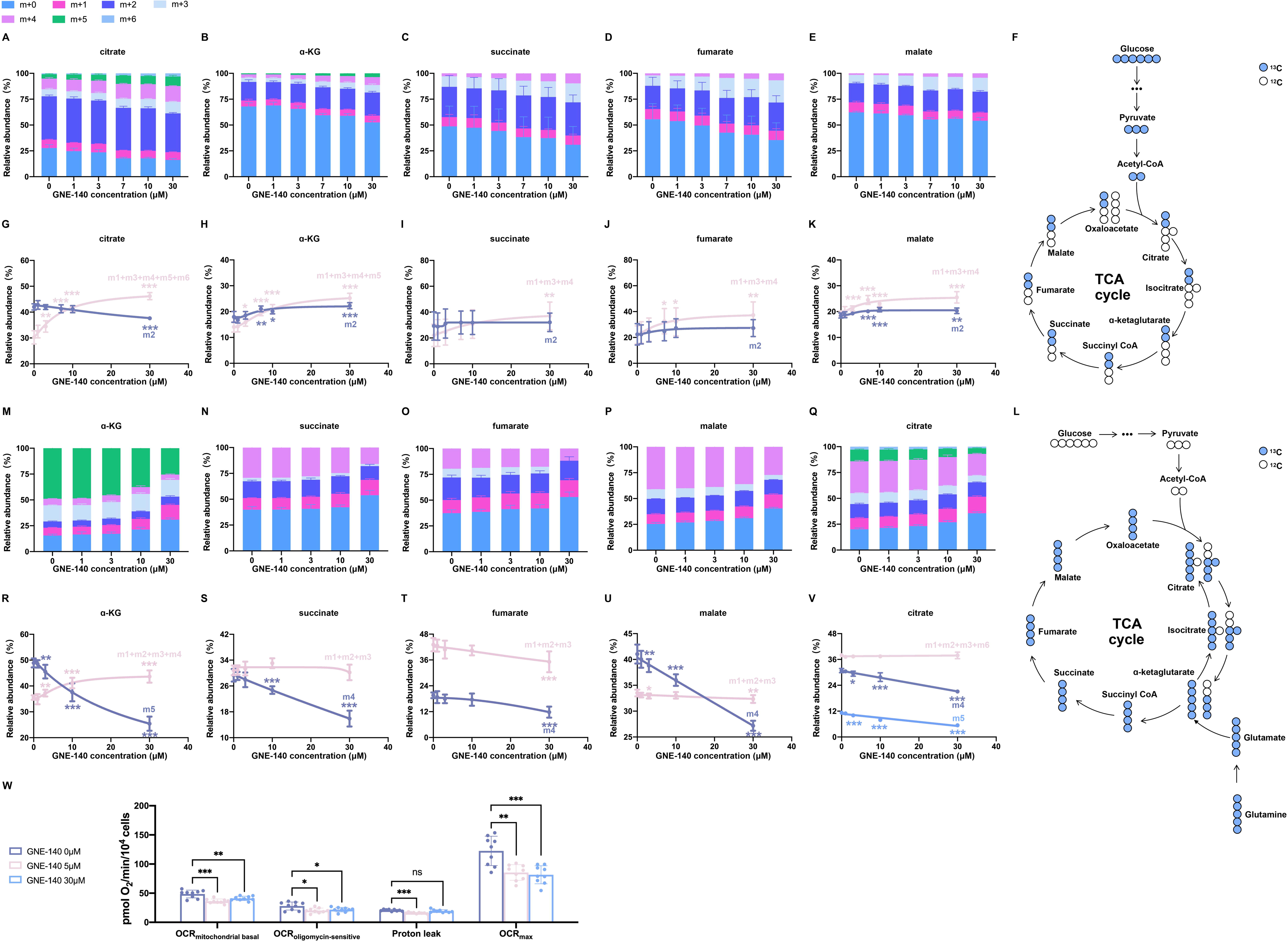
The effect GNE-140 on TCA cycle and OXPHOS in HeLa-LDHBKO cells. (A-K) Tracing glucose carbon to TCA cycle intermediates (citrate, α-KG, succinate, fumarate, malate). Cells were cultured in complete RPMI-1640 medium containing 6 mM [^13^C_6_]glc with or without GNE-140 in a CO_2_ incubator for 6 hours, and then the percentages of isotopologues of the TCA cycle intermediates in cells were determined by LC-MS/MS as described in Materials and Methods. (A-E) The total ^13^C labeling of the TCA cycle intermediates. (F) A metabolic diagram of the isotope labeling of TCA cycle when [^13^C_6_]glc was used as labeling substrate. (G-K) The isotope labeling pattern of the TCA cycle intermediates, including m2 isotopologues% and the sum of other isotopologues% (m1 + m3 + m4 + m5 + m6 for citrate, m1 + m3 + m4 + m5 for α-KG, m1 + m3 + m4 for succinate/fumarate/malate). (L-V) Tracing glutamine carbon to TCA cycle intermediates (citrate, α-KG, succinate, fumarate, malate). Cells were cultured in complete RPMI-1640 medium containing 2 mM [^13^C_5_]gln with or without GNE-140 in a CO_2_ incubator for 6 hours, and then the percentages of isotopologues of the TCA cycle intermediates in cells were determined by LC-MS/MS as described in Materials and Methods. (L) A metabolic diagram of the isotope labeling of TCA cycle when [^13^C_5_]gln was used as labeling substrate. (M-Q) The total ^13^C labeling of the TCA cycle intermediates. (R-V) The isotope labeling pattern of the TCA cycle intermediates, including m5 α-KG%, m4 succinate%, m4 fumarate%, m4 malate%, m4 citrate%, m5 citrate%, and the sum of other isotopologues% (m1 + m2 + m3 + m4 for α-KG, m1 + m2 + m3 for succinate/fumarate/malate, m1 + m2 + m3 + m6 for citrate). (W) OCR with or without GNE-140, measured as described in Materials and Methods. Data are mean ± SD from 3 independent experiments. *, *P*<0.05, **, *P*<0.01, ***, *P*<0.001.

To resolve the contradictory results, we traced [^13^C_5_]glutamine into TCA cycle. Glutamine is a major source for replenishing TCA cycle intermediates through two deamination reactions (Figure 5L). When cells were incubated with GNE-140, the percentage of m5 α-KG decreased with increasing concentrations of GNE-140, while isotopologues% (m1, m2, m3, m4) significantly increased (Figure 5M & R). This decrease in m5 α-KG indicated reduced direct influx of labeled glutamine into the TCA cycle, while the increase in other α-KG isotopologues% suggested enhanced cycling of TCA cycle intermediates or reduced flux of intermediates leaving the TCA cycle. Moreover, m4 succinate, m4 fumarate, and m4 malate, directly derived from m5 α-KG, showed a significant decrease in cells incubated with GNE-140, while the percentage of other isotopologues (m1, m2, m3) remained unchanged (succinate and malate) or moderately decreased (fumarate) (Figure 5N-P, S-U). For citrate isotopologues, m4-citrate was directly from α-KG in the forward direction, while m5-citrate was likely generated through reductive carboxylation (Corbet & Feron, 2015; Leonardi, Subramanian, Jackowski, & Rock, 2012). In the presence of GNE-140, the percentages of m4-citrate and m5-citrate decreased significantly, while the percentage of other isotopologues (m1, m2, m3, and m6) remained unchanged (Figure 5Q & V). Tracing glutamine carbon into TCA cycle supported that GNE-140 reduced both glutamine carbon entrance into TCA cycle and leaving TCA cycle.

Combining the tracing data of [^13^C_6_]glucose and [^13^C_5_]glutamine, we could interpret why in [^13^C_6_]glucose tracing, the m2 α-KG increased with the increased concentration of GNE-140. It was because glutamine-carbon entrance into TCA cycle decreased with the increased concentration of GNE-140, consequently, the percentage of m2 α-KG increased with the increased concentration of GNE-140. This interpretation could be extended to glucose carbon labeling pattern of other TCA cycle intermediates (Figure 5C-E).

Integrated analysis of dual isotope tracing data demonstrates that LDH inhibition reduces both influx and efflux of the TCA cycle. Tracing glucose and glutamine carbon into TCA cycle supports that LDH inhibition causes reduced flux of glucose-derived acetyl CoA into TCA cycle, coupling with a reduced flux of glutamine-derived α-KG into TCA cycle, and a decrease in the flux of intermediates leaving TCA cycle. The results are compatible with theoretical prediction. Under any circumstance, the reactions by which TCA cycle intermediates distribute to other pathways and those by which they are replenished must be balanced. In GNE-140 group, glutamine carbon entering into TCA cycle was reduced, glucose carbon (acetyl CoA) entering into TCA cycle should be also reduced, or vice versa.

We then measured mitochondrial respiration in these cells. The mitochondrial oxygen consumption of HeLa-LDHBKO in the absence of GNE-140 was significantly higher than that in the presence of GNE-140 (Figure 5W). Notably, with GNE-140 treatment, the basal respiration, ATP associated oxygen consumption, and maximal respiration all decreased, indicating a reduced electron flux rate through ETC. Because mitochondrial respiration depends on the electron generated from TCA cycle, the reduced mitochondrial respiration also meant the reduced TCA cycle activity. By combining all the data of glucose tracing, glutamine tracing, and oxygen consumption measurement, we conclude that LDH inhibition leads to reduced TCA cycle and impaired OXPHOS.

We repeated the above experiments on HeLa-Ctrl and HeLa-LDHAKO cells, including isotope tracing of TCA cycle intermediates using [^13^C_6_]glucose (Figure 5 — figure supplement 1)and [^13^C_5_] glutamine (Figure 5 — figure supplement 2), oxygen consumption rate (Figure 5—figure supplement 3). The results showed that GNE-140 inhibited both TCA cycle and OXPHOS. The effect of GNE-140 on the labeling pattern of TCA cycle intermediates by [^13^C_6_]glucose and [^13^C_5_]glutamine in HeLa-Ctrl and HeLa-LDHAKO were similar to that in HeLa-LDHBKO cells. The inhibition extent of mitochonidrial respiration by GNE-140 from high to low was HeLa-LDHBKO, HeLa-LDHAKO, and HeLa-Ctrl, which was agreeable with the inhibition extent of LDH in these cells, the residual LDH activity from low to high in the presence of GNE-140 was HeLa-LDHBKO, HeLa-LDHAKO, and HeLa-Ctrl (Figure 5—figure supplement 4). It was noted that GNE-140 did not significantly inhibit OXPHOS in HeLa-Ctrl, the possible reason was explained in the section of Discussion. LDHA KO combined with GNE-140 also inhibited the mitochondrial respiration. GNE-140 could also inhibit the mitochondrial respiration of 4T1-LDHAKO cells (Figure 5—figure supplement 5).

### The effect of GNE-140 on energy production, redox state, and cell survival

We observed that GNE-140 under normoxia significantly inhibited both glycolysis and mitochondrial respiration in HeLa-LDHBKO cells, and this agent under hypoxia was more efficacious in suppressing glycolysis, causing a marked reduction in the concentrations of ATP (Table supplement 9 & 10). These results prompted us to further analyze the energy metabolism and redox state in HeLa-LDHBKO cells responding to LDH inhibition by GNE-140.

We calculated the ATP production rate of the cells according to the glycolytic rate and mitochondrial respiration (Figure 6A). Under normoxia (21% oxygen), GNE-140 (30 μM) simultaneously inhibited glycolysis and mitochondrial respiration, resulting in a 58% decrease of total ATP output. Under hypoxia (1% oxygen), assuming OXPHOS rate was negligible, GNE-140 inhibited glycolysis-derived ATP (which was approximately to total ATP output) by 88%. Under normoxia, in the presence of GNE-140, ATP concentration reduced by 56%, ADP concentration remained unchanged, AMP concentration increased by 157%, and the pool size of AMP + ADP + ATP decreased by 50% (Figure 6B). Under hypoxia and in the presence of GNE-140, ATP and ADP concentrations decreased by 85% and 61%, respectively, AMP concentration did not change significantly, and the pooled concentration (AMP + ADP + ATP) decreased by 78% (Figure 6B). Together, GNE-140 not only markedly inhibited the ATP generation rate but also significantly reduced the pooled concentration of ATP + ADP + AMP.

**Figure 6.**
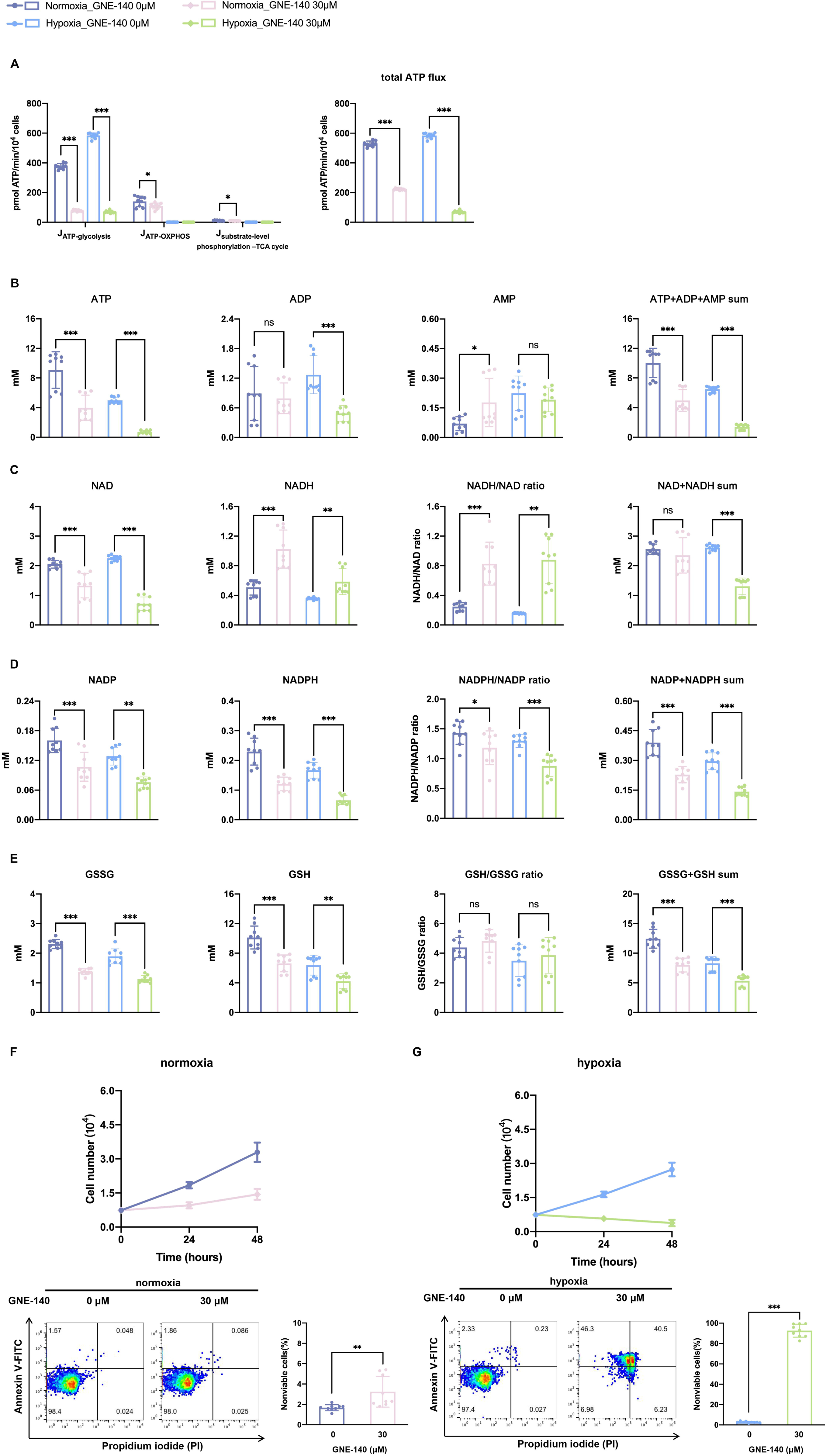
The effect of GNE-140 on energy production, redox state, and cell survival in HeLa-LDHBKO cells under normoxia and hypoxia. (A) ATP generation rate from glycolysis, OXPHOS, and substrate-level phosphorylation in TCA cycle, according to the methods described in Materials and Methods. Under hypoxia, we assume OXPHOS is negligible. (B) Cellular concentrations of ATP, ADP, and AMP (Table supplement 9 & 10). (C) Cellular concentrations of NAD^+^ and NADH (Table supplement 9 & 10). (D) Cellular concentrations of NADP^+^ and NADPH. (E) Cellular concentrations of GSH and GSSG. (F) Cell growth curves and cell death assays under normoxia. (G) Cell growth curves and cell death assays under hypoxia. Data are mean ± SD from 3 independent experiments. *, *P*<0.05, **, *P*<0.01, ***, *P*<0.001.

Two pairs of coenzymes (NAD^+^ and NADH, NADP^+^ and NADPH) could largely reflect the intracellular redox state. NAD^+^ and NADH are the major intermediate acceptors for transferring electrons in cellular catabolism. Under normoxia, after GNE-140 treatment, the concentration of NAD^+^ decreased and the concentration of NADH increased, resulting in an increased NADH/NAD^+^, but the pooled concentrations (NAD^+^ + NADH) did not change significantly (Figure 6C). Under hypoxia and with or without GNE-140, NAD^+^ concentrations were 2.26 mM and 0.72 mM, respectively, while NADH concentrations were 0.36 mM and 0.59 mM, respectively, and as a result, NADH/NAD^+^ increased by 4.5 folds but the pooled concentration (NADH + NAD^+^) decreased by 50 % (Figure 6C). Since NADP^+^ is derived from NAD^+^ via phosphorylation by NAD^+^ kinase, we speculated that the decrease in the total concentration (NADH + NAD^+^) could affect NADP^+^ and NADPH, which were intermediate acceptors that transfer electrons in cellular anabolism. Under normoxia or hypoxia, GNE-140 induced a decrease in the concentrations of NADP^+^ and NADPH (Figure 6D).

GSH is the most abundant anti-oxidant in cells. Oxidation of GSH converts to GSSG, which is recycled back to GSH via glutathione reductase using NADPH as the electron donor. It is well documented that a change in the concentration of NADPH or NADPH/NADP^+^ is directly associated with the concentration of GSH, GSSG, and/or GSH/GSSG (Ghosh, Levault, & Brewer, 2014; Xiao & Loscalzo, 2020; Zhou et al., 2016). Treatment of cells with GNE-140 under normoxia or hypoxia did not significantly affect GSH/GSSG, but did reduce the concentrations of both GSH and GSSG (Figure 6E).

The above results demonstrated that GNE-140 through inhibition of LDH could significantly alter the energy metabolism and redox state in cells, and that the inhibition was more efficacious under hypoxia than under normoxia. This differential inhibition under 2 conditions may lead to different effect on cell growth and survival. Indeed, GNE-140 (30 μM) under normoxia inhibited the growth of HeLa-LDHBKO cells but only exhibited marginally significant cytotoxicity (Figure 6F). In contrast, GNE-140 (30 μM) effectively killed HeLa-LDHBKO cells under hypoxia (Figure 6G).

Unlike HeLa-LDHBKO cells (Figure 6F & G), 30 μM GNE-140 moderately inhibited the growth of HeLa-Ctrl or HeLa-LDHAKO cells, but did not induce cell death (Figure 6—figure supplement 1). This was consistent with the potency of GNE-140, which was a much stronger inhibitor to LDHA than to LDHB (Figure 2A & B).

## Discussion

In contrast to prior findings suggesting that LDH inhibition results in the suppression of glycolysis while enhancing mitochondrial respiration, our study demonstrates that LDH inhibition leads to the suppression of not only glycolysis but also TCA cycle and OXPHOS. Understanding this inconsistency requires a mechanistic explanation. It is worth noting that previous publications have largely left the biochemical mechanism by which LDH regulates glycolysis, TCA cycle, and OXPHOS unexplored. Here, we provide a detailed examination of the kinetic and thermodynamic insights into how LDH regulates these biochemical processes, as discussed further below.

The regulation of the glycolytic pathway by LDH involves a dynamic interplay between enzyme kinetics and thermodynamic potentials within the pathway. Inhibition of LDH initiates a cascade of sequential kinetic and thermodynamic changes within glycolysis. Initially, the loss of LDH activity leads to an accumulation of free NADH and a depletion of free NAD^+^. The decrease in the free NAD^+^ concentration perturbs the activity of GAPDH, disrupting the balance in the concentrations of glycolytic intermediates, characterized by significant increases in the concentrations of GA3P, DHAP, FBP, and significant decrease in the concentration of 3PG. This disruption of glycolytic intermediates further changes the thermodynamic state of the reactions catalyzed by PFK1, aldolase, TPI, and PGAM, rendering these reactions significantly more favorable in the reverse direction. The combined kinetic and thermodynamic changes induced by inhibition of LDH contribute to the suppression of glycolysis. Based on our findings, the suppression of glycolysis through LDH inhibition primarily affects the segment encompassing PFK1, aldolase, TPI, GAPDH, PGK1, and PGAM.

The elucidation of the mechanistic insight into regulation of glycolysis by LDH has a broader significance in metabolic regulation. In literature and biochemistry textbooks, metabolic regulation is understood through two fundamental principles: chemical thermodynamics and enzyme kinetics. Chemical thermodynamics and enzyme kinetics each serve distinct roles: while the thermodynamics dictates the energetic favorability of a reaction or a metabolic pathway, the enzyme kinetics controls the rate. The glycolytic pathway, the most thoroughly studied metabolic pathway, serves as a prototypical model for metabolic regulation. The rate regulation of glycolysis is emphasized by the enzyme kinetics, including feedback regulation, feedforward activation, allosteric regulation, expression of enzymes, posttranslation modification of enzymes, etc, while the thermodynamic part focuses on Gibbs free energy, that determines the directionality and energy transfer in the pathway. However, the interaction between enzyme kinetics and the thermodynamic properties of the glycolytic pathway is often overlooked in the glycolytic regulation. In this study, we propose a model suggesting that glycolysis regulation occurs through the interaction between enzyme kinetics and the thermodynamic properties of the pathway. Therefore, our findings have implications that could contribute to a more comprehensive understanding of metabolic regulation in general.

It is essential to distinguish the roles of enzyme kinetics and the thermodynamic properties in isolated reactions from those in a metabolic pathway composed of multiple sequential reactions. Our study offers a model for this distinction. At steady state, the velocity of LDH is determined by the total activity of LDH and its substrate concentration. When the substrate concentration is fixed, the rate is proportional to the total activity, while the direction of the reaction is determined by the Gibbs free energy. However, in steady-state glycolysis, the rate of the LDH-catalyzed reaction is not directly proportional to its total activity. Inhibition of LDH induces serial kinetic and thermodynamic changes in the upper segment of the glycolytic pathway beyond the LDH’s catalytic step alone, ultimately influencing the glycolytic rate.

As shown in the result section, the inhibition of LDH decreases the rates of pyruvate production via glycolysis without significantly augmenting the proportion of pyruvate entering the mitochondria. Consequently, the flux of glycolysis-derived pyruvate to mitochondrial metabolism is diminished. This challenges the prior perception that LDH inhibition redirects pyruvate flux toward mitochondrial metabolism.

While LDH inhibition decreases the flux of glucose and glutamine carbon into TCA cycle, it also coincides with a reduction in the flux of intermediates exiting TCA cycle. As such, the extent of OXPHOS inhibition can vary depending on the ratios of these two flux rates, as observed in HeLa-LDHBKO, HeLa-LDHAKO, and HeLa-Ctrl cells.

The elucidated mechanism above explains why LDH inhibition results in the suppression of glycolysis, TCA cycle, and mitochondrial respiration. This relationship is vividly illustrated in the schematic diagram (Figure 7).

**Figure 7.**
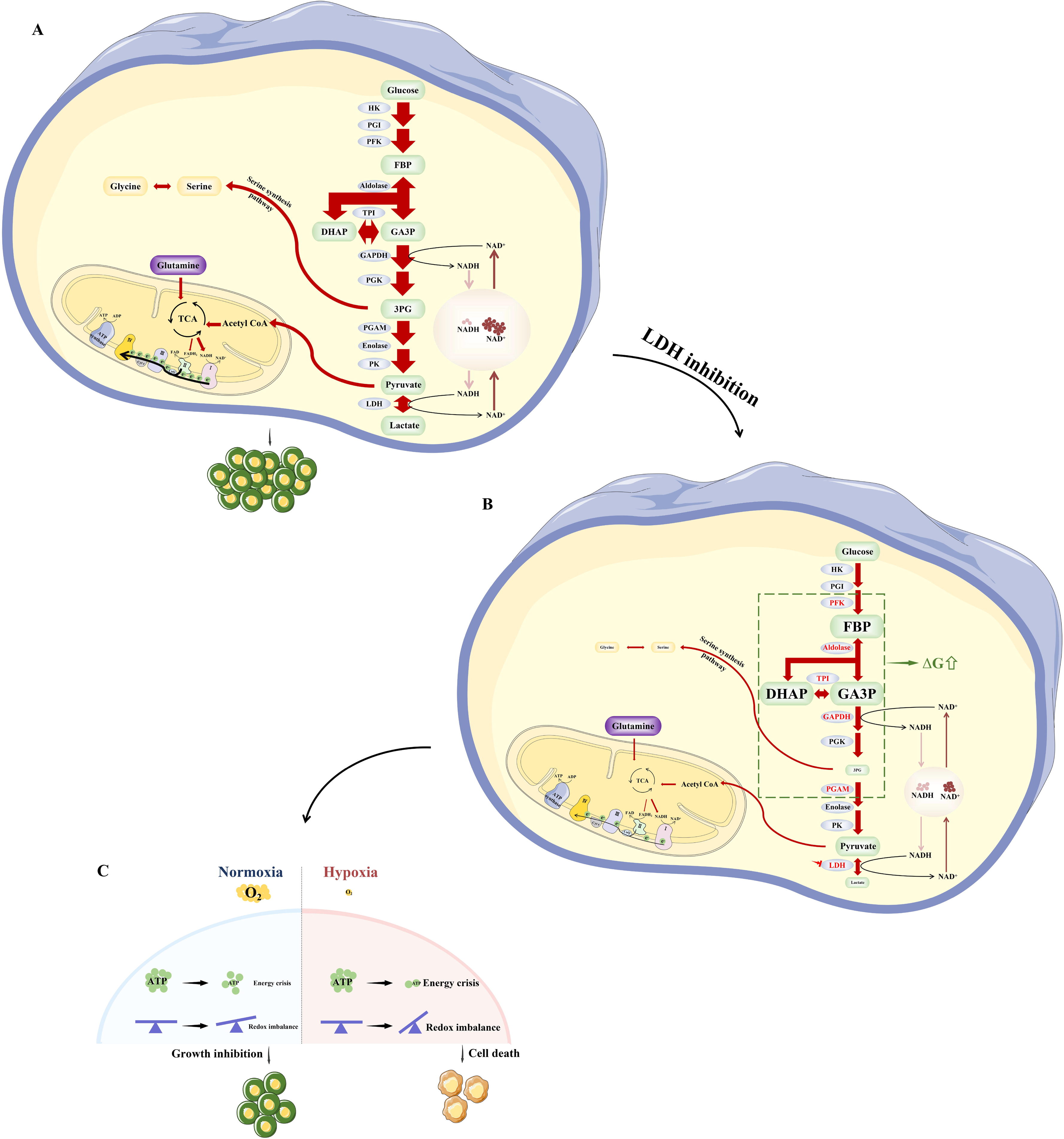
Schematic diagrams to demonstrate the impact on glycolysis, TCA cycle, and OXPHOS via regulation of LDH. (A) Tumor cells consume glucose through glycolysis. A significant portion of the pyruvate generated during glycolysis is converted into lactate, while a fraction enters the TCA cycle. This metabolic pathway aids in the production of electron donors such as NADH and FADH_2_, subsequently participating in OXPHOS. (B) When LDH is inhibited, it triggers a series of biochemical changes in the glycolytic pathway, resulting in the inhibition of glycolysis: the concentration of free NADH increases while the concentration of free NAD^+^ decreases; the decreased concentration of free NAD^+^ perturbs the activity of GAPDH, disrupting the balance of glycolytic intermediates, causing a marked increase of the concentrations in FBP, DHAP, GA3P, and significant decrease of the concentration in 3PG, leading to significant changes of ΔGs of the reactions catalyzed by PFK1, aldolase, TPI, and PGAM in the glycolytic pathway, favoring the reverse direction of these reactions. As the overall rate of pyruvate production decreases without significant proportion of pyruvate redirecting to mitochondrial metabolism, the flux of glucose carbon into the TCA cycle decreases, and the anaplerosis of glutamine to TCA cycle intermediates also decreases. Consequently, the TCA cycle and OXPHOS are inhibited. (C) Inhibiting LDH causes energy crisis and redox imbalance, particularly accentuated under hypoxia versus normoxia. As a result, inhibition of LDH activity inhibits growth under normoxia while kills cancer cells under hypoxia.

We observed that LDH activity correlated with the pooled concentrations of key variables (ATP + ADP + AMP, NAD^+^ + NADH, NADP^+^ + NADPH, GSH + GSSG) in cells. Deletion of LDH isoforms (LDHAKO or LDHBKO) did not significantly alter the concentrations of these variables. However, treatment of HeLa-LDHBKO cells with GNE-140 resulted in decreased concentrations of these variables. Considering the central roles of glycolysis, TCA cycle, and OXPHOS in cellular metabolism, which bridge catabolism and anabolism, changes in the synthesis and breakdown of these variables could be anticipated. Further investigation is required to elucidate how LDH influences the balance between the synthesis and breakdown of these variables by regulating glycolysis and mitochondrial respiration.

Inhibition of LDH induces energy crises and redox imbalances in cancer cells, with a much more profound impact under hypoxia than under normoxia condition. This difference is linked to distinct effects on cancer cell growth and death: LDH inhibition impairs cancer cell growth but does not induce cell death under normoxia, whereas it effectively kills cancer cells under hypoxia. As inhibition of LDH may effectively kill cancer cells under hypoxia, it may have potential for cancer treatment in the future, given that hypoxia confers cancer cells with resistance to radiation (Moeller et al., 2005; Mukherjee et al., 2009; Telarovic, Wenger, & Pruschy, 2021), immune checkpoint inhibitors (Chouaib, Noman, Kosmatopoulos, & Curran, 2017; Ding et al., 2021; Semenza, 2012), as well as numerous chemotherapeutic agents (Skarsgard, Vinczan, Skwarchuk, & Chaplin, 1994; Wilson, Keng, & Sutherland, 1989; Yokoi & Fidler, 2004).

Isotope tracing is a common method applied in metabolic analysis across all fields of biology. The percentage of TCA cycle intermediates labeled by [^13^C_6_]glucose is often interpreted as the efficiency of glucose carbon entering the TCA cycle. A higher percentage suggests higher efficiency, and vice versa. However, caution should be exercised when interpreting isotope tracing data. In this study, treatment of cells with GNE-140 led to an increase labeling percentage of TCA cycle intermediates by [^13^C_6_]glucose (Figure 5A-E). However, this does not necessarily imply an increase in glucose carbon flux into TCA cycle; rather, it indicates a reduction in both the flux of glucose carbon into TCA cycle and the flux of intermediates leaving TCA cycle. When interpreting the data, multiple factors must be considered, including the carbon-13 labeling pattern of the intermediates (m1, m2, m3, ---) (Figure 5G-K), replenishment of intermediates by glutamine (Figure 5M-V), and mitochondrial oxygen consumption rate (Figure 5W). All these factors should be taken into account to derive a proper interpretation of the data.

### Several additional issues deserve further discussion

#### The temporal and steady state metabolic regulation by LDH

Although very early time points (seconds to minutes) are essential for isolating the most proximal biochemical consequences of acute LDH inhibition before downstream propagation, the objective of the present study is different. Here, we focus on the metabolic steady state re-established after sustained suppression of LDH activity, because this adapted state is more informative for understanding long-term metabolic consequences and for predicting therapeutic outcomes of LDH-targeted interventions in cancer cells. In highly interconnected metabolic networks, transient changes immediately after perturbation often reflect short-lived imbalances, whereas the reconstructed steady state reflects the integrated constraints of redox balance, enzyme kinetics, and pathway thermodynamics that ultimately determine persistent flux behavior.

A similar consideration applies to the LDHA/LDHB knockout models. We recognize that generating stable knockout lines necessarily involves temporal perturbation and cellular adaptation. Nevertheless, our comparisons are explicitly made between two steady states: the baseline steady state of control cells and the adapted steady state achieved after stable genetic disruption of LDHA or LDHB. Notably, LDHA or LDHB knockout alone produced minimal changes in glycolytic flux and mitochondrial respiration, indicating that partial reduction of LDH activity can be compensated at steady state, consistent with the exceptionally high catalytic capacity of LDH relative to upstream glycolytic enzymes (Table supplement 1).

Importantly, our conclusions do not rely on a single inhibitor condition at a single time point. Instead, we established LDH-activity-dependent quantitative relationships linking residual LDH activity to pathway behavior over a broad inhibition range, providing strong evidence that the system occupies stable metabolic states at different levels of LDH activity rather than undergoing non-specific collapse (Figure 8). When LDH activity was reduced from 100% to approximately ∼9% — achieved by genetic perturbation and/or partial pharmacologic inhibition — glucose consumption and lactate production remained essentially unchanged, supporting maintenance of steady-state glycolytic flux despite substantial LDH suppression (Figure 8A & B). Only when LDH activity was further reduced below this threshold did glycolytic flux decline progressively, consistent with a nonlinear control structure in which LDH exerted limited control until its activity fell beneath a critical level.

**Figure 8.**
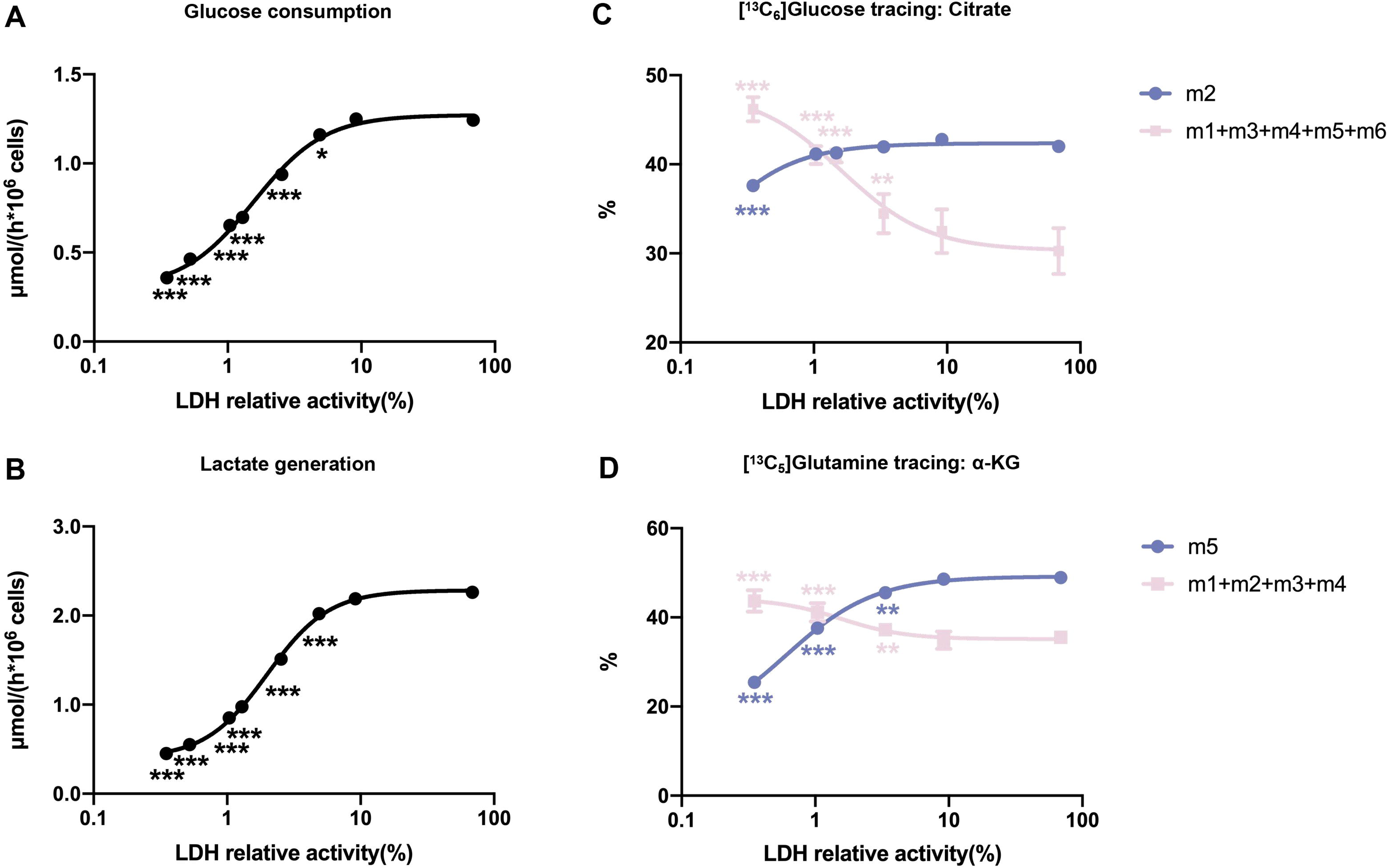
Quantitative relationship between LDH activity, glycolytic flux, and glucose-derived carbon entry into the TCA cycle in HeLa-LDHBKO cells. (A & B) Correlation between residual LDH activity and glycolytic flux (glucose consumption and lactate generation rates). The curves were generated by integrating glycolytic flux data (Figure 2C) with enzyme activity measurements from LDH inhibition using GNE-140 (Figure 2D). (C) LDH activity-dependent isotopic labeling patterns of citrate from [^13^C_6_]glucose. The m2 isotopologue percentage (reflecting initial glucose-derived acetyl-CoA entry into the TCA cycle) and the combined percentage of higher-order isotopologues (reflecting carbon retention and cycling) are plotted against LDH activity, based on isotopologue distributions (Figure 5G) and LDH activity levels (Figure 2D). (D) LDH activity-dependent isotopic labeling patterns of α-KG from [^13^C_5_]glutamine. The m5 isotopologue percentage (direct glutamine-derived α-KG) and the combined percentage of other isotopologues are plotted against LDH activity, using data from Figure 5R and Figure 2D. In all panels, LDH activity in HeLa-Ctrl cells is normalized to 100%. Data are presented as mean ± SD from 3 independent experiments. *, P < 0.05; **, P < 0.01; ***, P < 0.001.

Consistent with this threshold behavior, isotope tracing revealed distinct LDH-activity-dependent transitions in mitochondrial carbon utilization (Figure 8 C & D). Across the range from 100% to ∼9% LDH activity, the [^13^C_6_]glucose-derived isotopologue distribution of citrate remained largely unchanged, whereas deeper inhibition reduced m2 citrate and increased higher-order citrate isotopologues, consistent with altered relative contributions of carbon entry versus cycling/retention within the TCA cycle. Similarly, [^13^C_5_]glutamine tracing showed that deeper LDH inhibition decreased the direct m5 contribution to citrate in a graded manner, accompanied by corresponding increases in other isotopologues. Together, these quantitative transitions support the concept that LDH regulates metabolism in a state-dependent manner, such that progressive loss of LDH activity drives the system into distinct steady states characterized by coordinated remodeling of glycolysis, the TCA cycle, and oxidative phosphorylation.

#### Reconciling discrepancies with prior studies

Multiple prior studies have reported that LDH inhibition in cancer cells was accompanied by increased oxygen consumption or enhanced oxidative metabolism. At first glance, these observations appear inconsistent with our findings that sufficiently strong LDH suppression can coordinately reduce glycolysis, TCA cycle activity, and OXPHOS. We note, however, that the prevailing expectation of a “switch” from fermentation to respiration following LDH inhibition is often framed by analogy to the classical Pasteur and Crabtree effects, and this analogy may be mechanistically misleading.

In the Pasteur effect, fermentation increases primarily because oxygen becomes limiting, i.e., the restriction of the terminal electron acceptor constrains mitochondrial electron transport and enforces reliance on anaerobic ATP production. In the classical Crabtree effect, high glucose availability suppresses respiration through regulatory mechanisms while glycolytic flux remains high, yielding aerobic fermentation even in the presence of oxygen. In both cases, the core determinant is the availability or regulation of mitochondrial respiratory capacity, rather than direct inhibition of a specific cytosolic enzymatic reaction.

By contrast, LDH inhibition constitutes a distinct biochemical perturbation: it directly disrupts cytosolic redox recycling by limiting NADH-to-NAD^+^ regeneration, thereby restricting NAD^+^-dependent steps in upper glycolysis (most notably GAPDH) and reshaping the thermodynamic state of near-equilibrium reactions within the pathway. Under conditions where LDH inhibition sufficiently reduces effective NAD^+^ availability and constrains glycolytic flux upstream, pyruvate generation decreases and carbon entry into the TCA cycle becomes limited, resulting in suppression of OXPHOS—consistent with our flux measurements, metabolite profiling, thermodynamic analyses, and isotope tracing. In this context, interpreting LDH inhibition as a Pasteur/Crabtree-like toggle may oversimplify the biochemical consequences of disrupting cytosolic NAD^+^ regeneration and its pathway-level kinetic–thermodynamic propagation.

#### The positive ΔG of aldolase, TPI, and PGAM in the glycolytic pathway

The positive ΔG of aldolase, TPI, and PGAM (Figure 3D & 4C) requires explanation. These reactions are tightly linked to both upstream and downstream reactions in the glycolytic pathway. In glycolysis, three key reactions catalyzed by HK2, PFK1, and PK are highly exergonic, providing the driving force for the conversion of glucose to pyruvate. The other reactions, including the one catalyzed by PGAM, operate near thermodynamic equilibrium and primarily serve to equilibrate glycolytic intermediates rather than control the overall direction of glycolysis, as previously described by us (Jin et al., 2024).

The endergonic nature of these reactions does not prevent it from proceeding in the forward direction. Instead, the directionality of the pathway is dictated by the exergonic reaction of PFK1 upstream, which pushes the flux forward, and by PK downstream, which pulls the flux through the pathway. The combined effects of PFK1 and PK may account for the observed endergonic state of these reactions.

However, if these reaction were isolated from the glycolytic pathway, it would tend toward equilibrium and never surpass it, as there would be no driving force to move the reaction forward.

## Materials and Methods

### Reagents

Reagents purchased from Sigma included: pyruvate (#V900232), NADH (#N8129), glucose (#G8270), lactic acid (#L1750), KCl (#P3911), Na_2_HPO_4_ (#S9763), MgCl_2_ (#442615), EDTA (#E9884), HEPES (#H3375), ATP (#A3377), NADP (#N8035), hexokinse (#H4502), glucose-6-phosphate dehydrogenase (G6PDH) (#G8404), glycine (#G7126), hydrazine (#H4766), LDH (#L2500), NAD (#N0632), NaOH (#S5881), NADPH (#N7505), HCl (#H9892), glucose-6-phosphate (G6P) (#G7879), pyridine (#P57506), EDC (#T511307), 3-NPH (#N21804), acetonitrile (#AX0156), formic acid (#F0507), K_2_CO_3_ (#209619), PGI (#P5381), α-GPDH (#G6751), TPI (#T6258), Aldolase (#A8811), PGK (#P7634), GAPDH (#G2267), ADP (#A5285), PK (#P7768), Enolase (#E6126), triethylamine (#TX1202), antimycin (#A8674), rotenone (#R8875), FCCP (#C2920), 2-vinylpyridine (#132292), triethanolamine (#T1377), DTNB (#D8130), GR (#G3664). Reagents purchased from MedChemExpress included: GNE-140 (#HY-100742), oligomycin (#HY-N6782). [^13^C_6_]D-Glucose (#CLM-1396-1) was purchased from Cambridge Isotope Laboratories. MTS/PES (#G3581) was purchased from Promega. AQC (#A131410) and methanol (#M116125) were purchased from Aladdin.

### LDH KO cells and cell cultures

LDHA KO cells, LDHB KO cells, and their control cells (HeLa-LDHAKO, HeLa-LDHBKO, HeLa-Ctrl, 4T1-LDHAKO, 4T1-LDHBKO, and 4T1-Ctrl cells) were established by us previously as reported(Wu et al., 2021; M. Ying, Guo, & Hu, 2019). Cells were all maintained in RPMI-1640 medium supplemented with 10% fetal bovine serum, 100 μg/ml penicillin/streptomycin in a humidified incubator at 37 in a 5% CO_2_ atmosphere. For culture under hypoxia, cells were placed in a hypoxia workstation (Xvivo System® Model X3 Cell Incubation and Handling Platform, BioSpherix, Ltd., RRID: SCR_021175) of 1% oxygen level when required.

### LDH enzyme activity assay and Ki calculation

The LDH enzyme activity was determined as described by us previously(Jin et al., 2020). Briefly, 1 mL reaction buffer containing substrates (25 μM-0.16 mM NADH and 2 mM pyruvate) was added to the cuvette of a spectrophotometer (DU® 700, Beckman Coulter). The reaction was started by adding cell lysate, and then absorbance at 340 nm was recorded. When determining the Ki value of the LDH inhibitor GNE-140, the procedure is the same as above except that the reaction mixture also contain GNE-140. With or without GNE-140, the enzyme activity curves responding to different NADH concentrations were obtained, and the Km and Ki value were calculated by Graphpad Prism software.

### Measurement of glucose and lactate

The concentrations of glucose and lactate were determined as described by us previously (Jin et al., 2020; Zeng & Hu, 2023; X. Zhu et al., 2021). Cells were plated in density of about 6×10^4^/well in 48-well plate overnight. After replacing with fresh culture medium for 6 hours (treated with different culture conditions, with GNE-140/without GNE-140/normoxia/hypoxia), medium was collected for following measurement. 10 μL samples or standard solution of glucose/ lactate and 190 μL reaction buffer were added into each well of 96-well plate, thoroughly mixed, and the absorbance at 340 nm was recorded using a multi-mode microplate reader (SpectraMax i3, Molecular Devices) 60 minutes after the reaction. The reaction buffer for glucose determination is composed of 100 mM KCl, 5 mM Na_2_HPO_4_, 5 mM MgCl_2_, 0.5 mM EDTA, 2 mM ATP, 0.2 mM NADP, 0.2 U/mL HK2, and 0.2 U/mL G6PDH, 200 mM Hepes, pH 7.4. The reaction buffer for lactate determination is composed of 200 mM glycine, 170 mM hydrazine, 2 mM NAD, and 5 U/mL LDH, pH 9.2.

### Measurement of intracellular glycolytic intermediates

Intracellular glycolytic intermediates (G6P, F6P, DHAP, GA3P, FBP, 3PG, Pyr, PEP, and 2PG) were measured as described previously(Jin et al., 2020; X. Zhu et al., 2021) with some modification. Cells were plated in density of about 3×10^6^/dish in 10cm-dishes overnight. After replacing with fresh culture medium for 6 hours (treated with different culture conditions, with GNE-140/without GNE-140/normoxia/hypoxia), cells were washed by pre-cold PBS twice, and added 600 μL 1 M pre-cold HClO_4_ to every 3 dishes of 10cm-dishes. Cells were collected by a scraper, kept on ice for 30 minutes, and neutralized by 100 μL 3 M K_2_CO_3_. Then, the mixture was centrifuged for 10 minutes (15,000 rpm/ 4 ). The reaction buffer contained 100 mM KCl, 5 mM Na_2_HPO_4_, 5 mM MgCl_2_, 0.5 mM EDTA, and 200 mM Hepes (pH 7.4).

G6P and F6P: 140 μL of supernatant and 0.2 mM NADP were added to 560 μL reaction buffer, and the reaction was started by adding 1 U/mL G6PDH. The first reaction to measure G6P ended when 340 nm absorbance reached a plateau, then 1 U/mL PGI was added to measure F6P.

FBP, DHAP and GA3P: 140 μL of supernatant and 0.1 mM NADH were added to 560 μL reaction buffer, the reaction to measure DHAP was started by adding 1 U/mL α-GPDH. When the first reaction ended, 1 U/ml TPI was added to measure GA3P. Finally, 1 U/ml Aldolase was added to measure FBP.

3PG: 140 μL of supernatant, 2 mM ATP, 0.1 mM NADH, 1 U/mL PGK were added to 560 μL reaction buffer, and the reaction was started by adding 1 U/ml GAPDH.

2PG, PEP and Pyr: 140 μL of supernatant, 0.1 mM NADH were added to 560 μl reaction buffer, the reaction to measure Pyr was started by adding 1 U/mL LDH. When the first reaction ended, 2 mM ADP and 1 U/mL PK were added to measure PEP. Finally, 1 U/mL Enolase was added to measure 2PG.

### Determination of intracellular ATP, ADP, AMP, NAD^+^, NADH, NADP^+^, NADPH, GSH, and GSSG

Cells were plated in density of about 5×10^5^/well in 6-well plate overnight. After replacing with fresh culture medium for 6 hours (treated with different culture conditions, with GNE-140/without GNE-140/normoxia/hypoxia), cells were washed with PBS three times and intracellular metabolites were extracted by adding 600 μL 80% pre-cold methanol and incubating in - 80 for 20 minutes.

ATP, ADP, AMP, NAD^+^, NADH were determined as described by us previously(Jin et al., 2020; X. Zhu et al., 2021). The extract was collected and centrifuged for 10 minutes (15,000 rpm/ 4 ). The supernatant was evaporated by a vacuum centrifugal concentrator and dissolved in 100 μL water for following ultra performance liquid chromatography (UPLC) analysis. Waters ACQUITY UPLC system with an ACQUITY UPLC HSS T3 column was used to perform the liquid chromatography. Solvent A is composed of 1% acetonitrile, 99% water, 20 mM triethylamine, pH 6.5, and solvent B is composed of 100% acetonitrile. The solvent gradient program is: 0 min, 100% solvent A; 3 min, 100% solvent A; 4 min, 98.5% solvent A; 7 min, 92% solvent A; 7.1 min, 100% solvent A; 10 min, 100% solvent A. 10 μL sample or standard solution was injected to perform the analysis with a flow rate at 0.3 mL/min. During the performance, the column was kept at 40 .

NADP^+^ and NADPH were measured according to the methods described by us previously(M. Ying et al., 2021). The extract was collected and centrifuged for 10 minutes (15,000 rpm/ 4 ). The supernatant was equally divided into two tubes and evaporated by a vacuum centrifugal concentrator. One tube of the sample was dissolved in 20 μL 0.01 M NaOH to determine the NADPH concentration, and the other tube was dissolved in equal volume of 0.01 M HCl to determine NADP. The tubes were incubated at 60 for 15 minutes. 10 μL samples or standard solution and 190 μL reaction buffer were added into each well of 96-well plate, and the absorbance at 490 nm was recorded 30 minutes after the reaction at 37 . The reaction buffer was composed of 100 mM Tris-HCl (pH 8.1, with 5 mM EDTA), 2 mM G6P, 1 U/mL G6PDH, 10 μL MTS/PES.

GSSG and GSH were measured according to the methods described previously(M. Ying et al., 2021; X. Zhu et al., 2021). The extract was collected and centrifuged for 10 minutes (15,000 rpm/ 4 ). The supernatant was evaporated by a vacuum centrifugal concentrator and dissolved in 400 μl 0.1 M KPE buffer, then equally divided into two tubes. 4 μL 2-vinylpyridine was added in one tube (tube A) to derivatize GSH for 1 hour at room temperature. Then 12 μl triethanolamine was added to tube A to neutralize for 10 minutes. 2 mg DTNB in 3 mL KPE, 2 mg NADPH in 3 mL KPE, and 40 μL GR (250 U/mL) in 3 mL KPE were included in the reaction buffer. 20 μL sample/blank/standard were added to wells of 96-well plate, then 120 μL mixture (including DTNB and GR) was added to each well. After 30 seconds, 60 μL NADPH was added and well mixed, then the absorbance at 412 nm was recorded immediately and measurements were taken every 30 seconds for 2 minutes.

### Measurement of cytosolic free NADH/NAD^+^

Cytosolic free NAD^+^ and NADH redox states in living cells were determined using a highly responsive sensor SoNar developed by Yi Yang(Y. Zhao et al., 2015). Briefly, cells were transfected with plasmid SoNar and stable clones were selected. The stable cells were plated in density of about 2×10^5^/dish in glass bottom cell culture dishes (biosharp, BS-15-GJM) overnight. After replacing with fresh culture medium for 6 hours (treated with different culture conditions, with/without GNE-140), cells were observed under a laser confocal microscope (Zeiss LSM710, Carl Zeiss). Sets for microscope imaging: dual-excitation ratio imaging, 420-BP 30-nm or 480-BP 35-nm excitation, and 535-BP 40-nm emission. The SoNar sensor showed a distinct fluorescence response to NADH and NAD^+^. That is, the ratio of fluorescence intensities excited at 420 nm and 480 nm (F420 nm/F480 nm) was increased in the NADH-bound holo form of SoNar compared with the apo form of SoNar, whereas the ratio was decreased in the NAD^+^-bound holo form of SoNar compared with the apo form of SoNar(Y. Zhao et al., 2016). For fluorescence images of cells, raw data were exported to ImageJ for analysis. The dual-excitation ratio was determined pixel by pixel by dividing the 420-nm excitation image by the 480-nm excitation image of the same region. The data statistics of a single independent experiment included 3 different original fluorescence images for each treatment group, and the number of total counted cells was at least 200.

### Calculation of the Gibbs free energy change ΔG of glycolytic reactions

The ΔG was calculated as described by us previously(Jin et al., 2020). Briefly, ΔG was calculated according to the equation 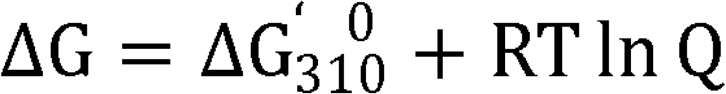, where 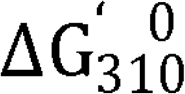 is the standard transformed Gibbs free energy at 37 and Q was calculated according to intermediate concentrations.

### Isotopic tracing by LC-MS/MS

Cells were plated in density of 5×10^5^/well in 6-well plate overnight. The cells were rinsed with PBS twice and cultured in RPMI-1640 containing 10% FBS and 6 mM [^13^C_6_] glucose for 6 hours (treated with different culture conditions, with/without GNE-140). Medium was collected for LC-MS/MS measurement. Meanwhile, cells were washed with PBS three times and intracellular metabolites were extracted by adding 600 μL 80% pre-cold methanol and incubating in - 80 for 20 minutes. The extract was collected and centrifuged for 10 minutes (15,000 rpm/ 4 ). The supernatant was evaporated by a vacuum centrifugal concentrator and dissolved in 50 μL water.

#### TCA cycle intermediates, pyruvate and lactate derivatization and LC-MS/MS method

The sample was derivatized according to previous study(Han, Gagnon, Eckle, & Borchers, 2013). Briefly, 10 μL sample (intracellular extract/medium) was mixed with sequentially with 10μL 7.5% pyridine in 75% methanol, 10 μL 150mM EDC in 100% methanol, 10μL 250mM 3-NPH in 50% methanol, and 10 μL 50% methanol. The reaction mechanism was the derivatization of carbonyl compounds to form 3-nitrophenylhydrazones. The mixture was incubated at room temperature for 60 minutes. An ACQUITY UPLC BEH C18 column attached to Waters ACQUITY UPLC system was used to perform liquid chromatography. Solvent A is composed of 0.1% formic acid, and solvent B is composed of 100% acetonitrile. The solvent gradient program is: 0 min, 80% solvent A; 6 min, 40% solvent A; 6.01 min, 0% solvent A; 7 min, 0% solvent A; 8 min, 80% solvent A; 10 min, 80% solvent A. 1 μL sample was injected to perform the analysis with a flow rate at 0.4 mL/min. During the performance, the column was kept at 55 .

#### Amino acid derivatization and LC-MS/MS method

The sample was derivatized according to the previous study(Salazar, Armenta, & Shulaev, 2012). Briefly, 10 μL sample (intracellular extract/medium) was added to 100 μL borate buffer (0.2 M borate buffer, pH 8.8), and mixed thoroughly. Then, 30 μL AQC (3 mg/mL) was added and vortexed immediately. The mixture was incubated at 60°C for 20 minutes. An ACCQ-TAG ULTRA C18 column attached to Waters ACQUITY UPLC system was used to perform liquid chromatography. Solvent A is composed of 1% acetonitrile, 99% water, 1.5 mM ammonia formate, pH 3.05, and solvent B is composed of 100% acetonitrile. The solvent gradient program is: 0 min, 80% solvent A; 6 min, 40% solvent A; 6.01 min, 0% solvent A; 7 min, 0% solvent A; 8 min, 80% solvent A; 10 min, 80% solvent A. 1 μL sample or standard solution was injected to perform the analysis with a flow rate at 0.7 mL/min. During the performance, the column was kept at 55 .

Isotopologues were analyzed by the mass spectrometer (X500 QTOF, Sciex). The MRM transitions (m/z), declustering potential, collision energy, and entrance potential were optimized for each metabolite (Table supplement 11 & 12).

### Measurement of mitochondrial respiration rate

Measurement of mitochondrial respiration rate was described by us previously (C. Ying, Jin, Zeng, Chao, & Hu, 2022; Zeng & Hu, 2023). Briefly, cells were plated in density of about 1×10^6^∼2×10^6^/dish in 6cm-dishes overnight. After replacing with fresh culture medium for 6 hours (treated with different culture conditions, with/without GNE-140), about 1×10^6^∼2×10^6^ cells were added to each O2K chamber (O2k-FluoRespirometer, ORBOROS), and the culture conditions in the chamber were the same as before (with/without GNE-140). Oligomycin (an inhibitor of ATP synthase, 2μg/mL), FCCP (an uncoupler of the electron transport and oxidative phosphorylation, 0.15-0.3μM, depending on different cells), rotenone (an inhibitor of respiratory complex I, 0.5μM), and antimycin A (an inhibitor of respiratory complex III, 2.5μM) were sequentially injected into chamber. OCR was calculated according to the following equations:

OCRmitochondrial basal = OCRcellular basal － OCRrot+aa; (OCRcellular basal is the oxygen consumption rate without the above mentioned inhibitors and uncoupler; OCR_rot+aa_ is the oxygen consumption rate after adding rotenone and antimycin A)

OCR_oligomycin-sensitive_ = OCR_cellular basal_ － OCR_oli_; (OCR_oli_ is the oxygen consumption rate after adding oligomycin)

Proton leak = OCR_oli_－ OCR_rot+aa_;

OCR_max_ =OCR_FCCP_ － OCR_rot+aa_; (OCR_FCCP_ is the oxygen consumption rate after adding FCCP)

OCR_non-mitochondrial_ = OCR_rot+aa_.

### Calculation of ATP generation rate

The ATP generation rate was calculated as described by us previously(Zeng & Hu, 2023). Briefly, the rate was calculated according to the equations as follows:

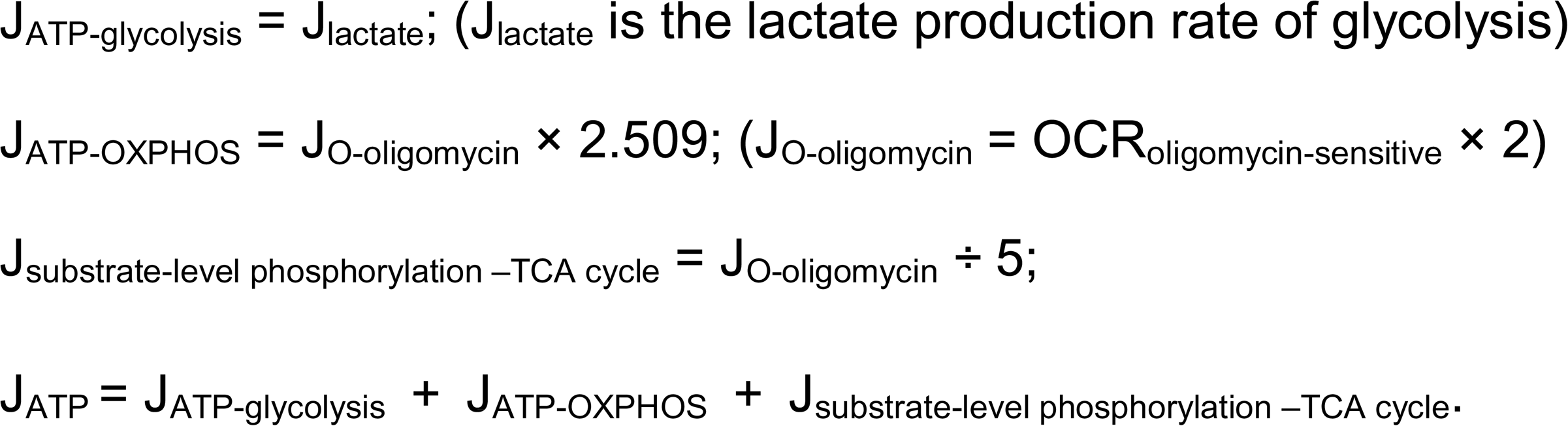

### Cell growth

Cell growth curves were obtained as described previously(C. Ying et al., 2022). Cells were plated in density of about 4×10^3^/well in 96-well plate overnight, and treated with different culture conditions (with GNE0140/without GNE-140/normoxia/hypoxia). Then, cells were counted at 0, 24, and 48 hours using a cell counter (BioTech, Countstar).

### Flow cytometric analysis

Flow cytometry was analyzed according to the methods described previously(C. Ying et al., 2022; Zeng & Hu, 2023). Cells were plated in density of about 3×10^4^/well, respectively, in 24-well plate overnight, and treated with different culture conditions (with GNE-140/without GNE-140/normoxia/hypoxia). After 48 hours, cells were collected, washed with PBS, stained with Annexin V-FITC (AV) and Propidium iodide (PI) solution (Beyotime, cat. no. C1062L) at room temperature in the dark for 10 minutes, and then the samples were analyzed by a flow cytometer (DxFLEX, Beckman Coulter).

### Statistics

All statistical analyses were performed using GraphPad Prism 8.0. The statistical significance of the difference between two groups was analyzed using unpaired two-tailed Student’s t-test. p < 0.05 was considered as statistically significant.

## Acknowledgments

We thank Professor Yi Yang (East China University of Science and Technology) for the kind gift of the SoNar sensor plasmid.

## Data availability

All data generated or analysed during this study are included in the manuscript and supporting files.

## References

Arseneault, R., Chien, A., Newington, J. T., Rappon, T., Harris, R., & Cumming, R. C. (2013). Attenuation of LDHA expression in cancer cells leads to redox-dependent alterations in cytoskeletal structure and cell migration. Cancer Lett, 338(2), 255–266. doi:10.1016/j.canlet.2013.03.034

Augoff, K., Hryniewicz-Jankowska, A., & Tabola, R. (2015). Lactate dehydrogenase 5: an old friend and a new hope in the war on cancer. Cancer Lett, 358(1), 1–7. doi:10.1016/j.canlet.2014.12.035

Boudreau, A., Purkey, H. E., Hitz, A., Robarge, K., Peterson, D., Labadie, S., . . . O’Brien, T. (2016). Metabolic plasticity underpins innate and acquired resistance to LDHA inhibition. Nat Chem Biol, 12(10), 779–786. doi:10.1038/nchembio.2143

Chen, Q., Xin, M., Wang, L., Li, L., Shen, Y., Geng, Y., . . . Fan, X. (2022). Inhibition of LDHA to induce eEF2 release enhances thrombocytopoiesis. Blood, 139(19), 2958–2971. doi:10.1182/blood.2022015620

Chen, X., Liu, L., Kang, S., Gnanaprakasam, J. R., & Wang, R. (2023). The lactate dehydrogenase (LDH) isoenzyme spectrum enables optimally controlling T cell glycolysis and differentiation. Sci Adv, 9(12), eadd9554. doi:10.1126/sciadv.add9554

Choi, S. Y., Collins, C. C., Gout, P. W., & Wang, Y. (2013). Cancer-generated lactic acid: a regulatory, immunosuppressive metabolite? J Pathol, 230(4), 350–355. doi:10.1002/path.4218

Chouaib, S., Noman, M. Z., Kosmatopoulos, K., & Curran, M. A. (2017). Hypoxic stress: obstacles and opportunities for innovative immunotherapy of cancer. Oncogene, 36(4), 439–445. doi:10.1038/onc.2016.225

Corbet, C., & Feron, O. (2015). Metabolic and mind shifts: from glucose to glutamine and acetate addictions in cancer. Curr Opin Clin Nutr Metab Care, 18(4), 346–353. doi:10.1097/MCO.0000000000000178

Cui, J., Shi, M., Xie, D., Wei, D., Jia, Z., Zheng, S., . . . Xie, K. (2014). FOXM1 promotes the warburg effect and pancreatic cancer progression via transactivation of LDHA expression. Clin Cancer Res, 20(10), 2595–2606. doi:10.1158/1078-0432.CCR-13-2407

Dantas, E., Erra Diaz, F., Pereyra Gerber, P., Merlotti, A., Varese, A., Ostrowski, M., . . . Geffner, J. (2016). Low pH impairs complement-dependent cytotoxicity against IgG-coated target cells. Oncotarget, 7(45), 74203–74216. doi:10.18632/oncotarget.12412

DeBerardinis, R. J., Mancuso, A., Daikhin, E., Nissim, I., Yudkoff, M., Wehrli, S., & Thompson, C. B. (2007). Beyond aerobic glycolysis: transformed cells can engage in glutamine metabolism that exceeds the requirement for protein and nucleotide synthesis. Proc Natl Acad Sci U S A, 104(49), 19345–19350. doi:10.1073/pnas.0709747104

Ding, X. C., Wang, L. L., Zhang, X. D., Xu, J. L., Li, P. F., Liang, H., . . . Hu, M. (2021). The relationship between expression of PD-L1 and HIF-1alpha in glioma cells under hypoxia. J Hematol Oncol, 14(1), 92. doi:10.1186/s13045-021-01102-5

Fantin, V. R., St-Pierre, J., & Leder, P. (2006). Attenuation of LDH-A expression uncovers a link between glycolysis, mitochondrial physiology, and tumor maintenance. Cancer Cell, 9(6), 425–434. doi:10.1016/j.ccr.2006.04.023

Fukumura, D., Xu, L., Chen, Y., Gohongi, T., Seed, B., & Jain, R. K. (2001). Hypoxia and acidosis independently up-regulate vascular endothelial growth factor transcription in brain tumors in vivo. Cancer Res, 61(16), 6020–6024.

Gatenby, R. A., & Gillies, R. J. (2004). Why do cancers have high aerobic glycolysis? Nat Rev Cancer, 4(11), 891–899. doi:10.1038/nrc1478

Ghosh, D., Levault, K. R., & Brewer, G. J. (2014). Relative importance of redox buffers GSH and NAD(P)H in age-related neurodegeneration and Alzheimer disease-like mouse neurons. Aging Cell, 13(4), 631–640. doi:10.1111/acel.12216

Han, J., Gagnon, S., Eckle, T., & Borchers, C. H. (2013). Metabolomic analysis of key central carbon metabolism carboxylic acids as their 3-nitrophenylhydrazones by UPLC/ESI-MS. Electrophoresis, 34(19), 2891–2900. doi:10.1002/elps.201200601

Jin, C., Hu, W., Wang, Y., Wu, H., Zeng, S., Ying, M., & Hu, X. (2024). Deciphering the interaction between PKM2 and the built-in thermodynamic properties of the glycolytic pathway in cancer cells. J Biol Chem, 300(9), 107648. doi:10.1016/j.jbc.2024.107648

Jin, C., Zhu, X., Wu, H., Wang, Y., & Hu, X. (2020). Perturbation of phosphoglycerate kinase 1 (PGK1) only marginally affects glycolysis in cancer cells. J Biol Chem, 295(19), 6425–6446. doi:10.1074/jbc.RA119.012312

Le, A., Cooper, C. R., Gouw, A. M., Dinavahi, R., Maitra, A., Deck, L. M., . . . Dang, C. V. (2010). Inhibition of lactate dehydrogenase A induces oxidative stress and inhibits tumor progression. Proc Natl Acad Sci U S A, 107(5), 2037–2042. doi:10.1073/pnas.0914433107

Leonardi, R., Subramanian, C., Jackowski, S., & Rock, C. O. (2012). Cancer-associated isocitrate dehydrogenase mutations inactivate NADPH-dependent reductive carboxylation. J Biol Chem, 287(18), 14615–14620. doi:10.1074/jbc.C112.353946

Li, X., Yang, Y., Zhang, B., Lin, X., Fu, X., An, Y., . . . Yu, T. (2022). Correction: Lactate metabolism in human health and disease. Signal Transduct Target Ther, 7(1), 372. doi:10.1038/s41392-022-01206-5

Liberti, M. V., & Locasale, J. W. (2016). The Warburg Effect: How Does it Benefit Cancer Cells? Trends Biochem Sci, 41(3), 211–218. doi:10.1016/j.tibs.2015.12.001

Maftouh, M., Avan, A., Sciarrillo, R., Granchi, C., Leon, L. G., Rani, R., . . . Giovannetti, E. (2014). Synergistic interaction of novel lactate dehydrogenase inhibitors with gemcitabine against pancreatic cancer cells in hypoxia. Br J Cancer, 110(1), 172–182. doi:10.1038/bjc.2013.681

Manerba, M., Vettraino, M., Fiume, L., Di Stefano, G., Sartini, A., Giacomini, E., . . . Recanatini, M. (2012). Galloflavin (CAS 568-80-9): a novel inhibitor of lactate dehydrogenase. ChemMedChem, 7(2), 311–317. doi:10.1002/cmdc.201100471

McCleland, M. L., Adler, A. S., Shang, Y., Hunsaker, T., Truong, T., Peterson, D., . . . Firestein, R. (2012). An integrated genomic screen identifies LDHB as an essential gene for triple-negative breast cancer. Cancer Res, 72(22), 5812–5823. doi:10.1158/0008-5472.CAN-12-1098

Moeller, B. J., Dreher, M. R., Rabbani, Z. N., Schroeder, T., Cao, Y., Li, C. Y., & Dewhirst, M. W. (2005). Pleiotropic effects of HIF-1 blockade on tumor radiosensitivity. Cancer Cell, 8(2), 99–110. doi:10.1016/j.ccr.2005.06.016

Mukherjee, B., McEllin, B., Camacho, C. V., Tomimatsu, N., Sirasanagandala, S., Nannepaga, S., . . . Burma, S. (2009). EGFRvIII and DNA double-strand break repair: a molecular mechanism for radioresistance in glioblastoma. Cancer Res, 69(10), 4252–4259. doi:10.1158/0008-5472.CAN-08-4853

Oshima, N., Ishida, R., Kishimoto, S., Beebe, K., Brender, J. R., Yamamoto, K., . . . Neckers, L. M. (2020). Dynamic Imaging of LDH Inhibition in Tumors Reveals Rapid In Vivo Metabolic Rewiring and Vulnerability to Combination Therapy. Cell Rep, 30(6), 1798–1810 e1794. doi:10.1016/j.celrep.2020.01.039

Park, J. O., Tanner, L. B., Wei, M. H., Khana, D. B., Jacobson, T. B., Zhang, Z., . . . Rabinowitz, J. D. (2019). Near-equilibrium glycolysis supports metabolic homeostasis and energy yield. Nat Chem Biol, 15(10), 1001–1008. doi:10.1038/s41589-019-0364-9

Peppicelli, S., Bianchini, F., & Calorini, L. (2014). Extracellular acidity, a "reappreciated" trait of tumor environment driving malignancy: perspectives in diagnosis and therapy. Cancer Metastasis Rev, 33(2-3), 823–832. doi:10.1007/s10555-014-9506-4

Perez-Escuredo, J., Dadhich, R. K., Dhup, S., Cacace, A., Van Hee, V. F., De Saedeleer, C. J., . . . Sonveaux, P. (2016). Lactate promotes glutamine uptake and metabolism in oxidative cancer cells. Cell Cycle, 15(1), 72–83. doi:10.1080/15384101.2015.1120930

Perez-Tomas, R., & Perez-Guillen, I. (2020). Lactate in the Tumor Microenvironment: An Essential Molecule in Cancer Progression and Treatment. Cancers (Basel), 12(11). doi:10.3390/cancers12113244

Petrelli, F., Cabiddu, M., Coinu, A., Borgonovo, K., Ghilardi, M., Lonati, V., & Barni, S. (2015). Prognostic role of lactate dehydrogenase in solid tumors: a systematic review and meta-analysis of 76 studies. Acta Oncol, 54(7), 961–970. doi:10.3109/0284186x.2015.1043026

Raghunand, N., Gatenby, R. A., & Gillies, R. J. (2003). Microenvironmental and cellular consequences of altered blood flow in tumours. Br J Radiol, 76 *Spec No 1*, S11-22. doi:10.1259/bjr/12913493

Rai, G., Brimacombe, K. R., Mott, B. T., Urban, D. J., Hu, X., Yang, S. M., . . . Maloney, D. J. (2017). Discovery and Optimization of Potent, Cell-Active Pyrazole-Based Inhibitors of Lactate Dehydrogenase (LDH). J Med Chem, 60(22), 9184–9204. doi:10.1021/acs.jmedchem.7b00941

Salazar, C., Armenta, J. M., & Shulaev, V. (2012). An UPLC-ESI-MS/MS Assay Using 6- Aminoquinolyl-N-Hydroxysuccinimidyl Carbamate Derivatization for Targeted Amino Acid Analysis: Application to Screening of Arabidopsis thaliana Mutants. Metabolites, 2(3), 398–428. doi:10.3390/metabo2030398

Semenza, G. L. (2012). Hypoxia-inducible factors in physiology and medicine. Cell, 148(3), 399–408. doi:10.1016/j.cell.2012.01.021

Skarsgard, L. D., Vinczan, A., Skwarchuk, M. W., & Chaplin, D. J. (1994). The effect of low pH and hypoxia on the cytotoxic effects of SR4233 and mitomycin C in vitro. Int J Radiat Oncol Biol Phys, 29(2), 363–367. doi:10.1016/0360-3016(94)90290-9

Telarovic, I., Wenger, R. H., & Pruschy, M. (2021). Interfering with Tumor Hypoxia for Radiotherapy Optimization. J Exp Clin Cancer Res, 40(1), 197. doi:10.1186/s13046-021-02000-x

Vander Heiden, M. G., Cantley, L. C., & Thompson, C. B. (2009). Understanding the Warburg effect: the metabolic requirements of cell proliferation. Science, 324(5930), 1029–1033. doi:10.1126/science.1160809

Walenta, S., Wetterling, M., Lehrke, M., Schwickert, G., Sundfor, K., Rofstad, E. K., & Mueller-Klieser, W. (2000). High lactate levels predict likelihood of metastases, tumor recurrence, and restricted patient survival in human cervical cancers. Cancer Res, 60(4), 916–921.

Warburg, O. (1956). On the origin of cancer cells. Science, 123(3191), 309–314. doi:10.1126/science.123.3191.309

Wilson, R. E., Keng, P. C., & Sutherland, R. M. (1989). Drug resistance in Chinese hamster ovary cells during recovery from severe hypoxia. J Natl Cancer Inst, 81(16), 1235–1240. doi:10.1093/jnci/81.16.1235

Wu, H., Wang, Y., Ying, M., Jin, C., Li, J., & Hu, X. (2021). Lactate dehydrogenases amplify reactive oxygen species in cancer cells in response to oxidative stimuli. Signal Transduct Target Ther, 6(1), 242. doi:10.1038/s41392-021-00595-3

Xiao, W., & Loscalzo, J. (2020). Metabolic Responses to Reductive Stress. Antioxid Redox Signal, 32(18), 1330–1347. doi:10.1089/ars.2019.7803

Xiao, W., Wang, R. S., Handy, D. E., & Loscalzo, J. (2018). NAD(H) and NADP(H) Redox Couples and Cellular Energy Metabolism. Antioxid Redox Signal, 28(3), 251–272. doi:10.1089/ars.2017.7216

Xie, H., Hanai, J., Ren, J. G., Kats, L., Burgess, K., Bhargava, P., . . . Seth, P. (2014). Targeting lactate dehydrogenase--a inhibits tumorigenesis and tumor progression in mouse models of lung cancer and impacts tumor-initiating cells. Cell Metab, 19(5), 795–809. doi:10.1016/j.cmet.2014.03.003

Xie, J., Dai, C., & Hu, X. (2016). Evidence That Does Not Support Pyruvate Kinase M2 (PKM2)- catalyzed Reaction as a Rate-limiting Step in Cancer Cell Glycolysis. J Biol Chem, 291(17), 8987–8999. doi:10.1074/jbc.M115.704825

Yeung, C., Gibson, A. E., Issaq, S. H., Oshima, N., Baumgart, J. T., Edessa, L. D., . . . Heske, C. M. (2019). Targeting Glycolysis through Inhibition of Lactate Dehydrogenase Impairs Tumor Growth in Preclinical Models of Ewing Sarcoma. Cancer Res, 79(19), 5060–5073. doi:10.1158/0008-5472.CAN-19-0217

Ying, C., Jin, C., Zeng, S., Chao, M., & Hu, X. (2022). Alkalization of cellular pH leads to cancer cell death by disrupting autophagy and mitochondrial function. Oncogene, 41(31), 3886–3897. doi:10.1038/s41388-022-02396-6

Ying, M., Guo, C., & Hu, X. (2019). The quantitative relationship between isotopic and net contributions of lactate and glucose to the tricarboxylic acid (TCA) cycle. J Biol Chem, 294(24), 9615–9630. doi:10.1074/jbc.RA119.007841

Ying, M., You, D., Zhu, X., Cai, L., Zeng, S., & Hu, X. (2021). Lactate and glutamine support NADPH generation in cancer cells under glucose deprived conditions. Redox Biol, 46, 102065. doi:10.1016/j.redox.2021.102065

Yokoi, K., & Fidler, I. J. (2004). Hypoxia increases resistance of human pancreatic cancer cells to apoptosis induced by gemcitabine. Clin Cancer Res, 10(7), 2299–2306. doi:10.1158/1078-0432.ccr-03-0488

Zdralevic, M., Brand, A., Di Ianni, L., Dettmer, K., Reinders, J., Singer, K., . . . Kreutz, M. (2018). Double genetic disruption of lactate dehydrogenases A and B is required to ablate the "Warburg effect" restricting tumor growth to oxidative metabolism. J Biol Chem, 293(41), 15947–15961. doi:10.1074/jbc.RA118.004180

Zeng, S., & Hu, X. (2023). Lactic acidosis switches cancer cells from dependence on glycolysis to OXPHOS and renders them highly sensitive to OXPHOS inhibitors. Biochem Biophys Res Commun, 671, 46–57. doi:10.1016/j.bbrc.2023.05.097

Zhao, Y., Hu, Q., Cheng, F., Su, N., Wang, A., Zou, Y., . . . Yang, Y. (2015). SoNar, a Highly Responsive NAD+/NADH Sensor, Allows High-Throughput Metabolic Screening of Anti-tumor Agents. Cell Metab, 21(5), 777–789. doi:10.1016/j.cmet.2015.04.009

Zhao, Y., Wang, A., Zou, Y., Su, N., Loscalzo, J., & Yang, Y. (2016). In vivo monitoring of cellular energy metabolism using SoNar, a highly responsive sensor for NAD(+)/NADH redox state. Nat Protoc, 11(8), 1345–1359. doi:10.1038/nprot.2016.074

Zhao, Z., Han, F., Yang, S., Wu, J., & Zhan, W. (2015). Oxamate-mediated inhibition of lactate dehydrogenase induces protective autophagy in gastric cancer cells: involvement of the Akt-mTOR signaling pathway. Cancer Lett, 358(1), 17–26. doi:10.1016/j.canlet.2014.11.046

Zhou, L., Wang, F., Sun, R., Chen, X., Zhang, M., Xu, Q., Ye, D. (2016). SIRT5 promotes IDH2 desuccinylation and G6PD deglutarylation to enhance cellular antioxidant defense. EMBO Rep, 17(6), 811–822. doi:10.15252/embr.201541643

Zhu, X. H., Lu, M., Lee, B. Y., Ugurbil, K., & Chen, W. (2015). In vivo NAD assay reveals the intracellular NAD contents and redox state in healthy human brain and their age dependences. Proc Natl Acad Sci U S A, 112(9), 2876–2881. doi:10.1073/pnas.1417921112

Zhu, X., Jin, C., Pan, Q., & Hu, X. (2021). Determining the quantitative relationship between glycolysis and GAPDH in cancer cells exhibiting the Warburg effect. J Biol Chem, 296, 100369. doi:10.1016/j.jbc.2021.100369

