## Supplementary figures for "Elucidating the kinetic and thermodynamic insight into regulation of glycolysis by lactate dehydrogenase and its impact on tricarboxylic acid cycle and oxidative phosphorylation in cancer cells"

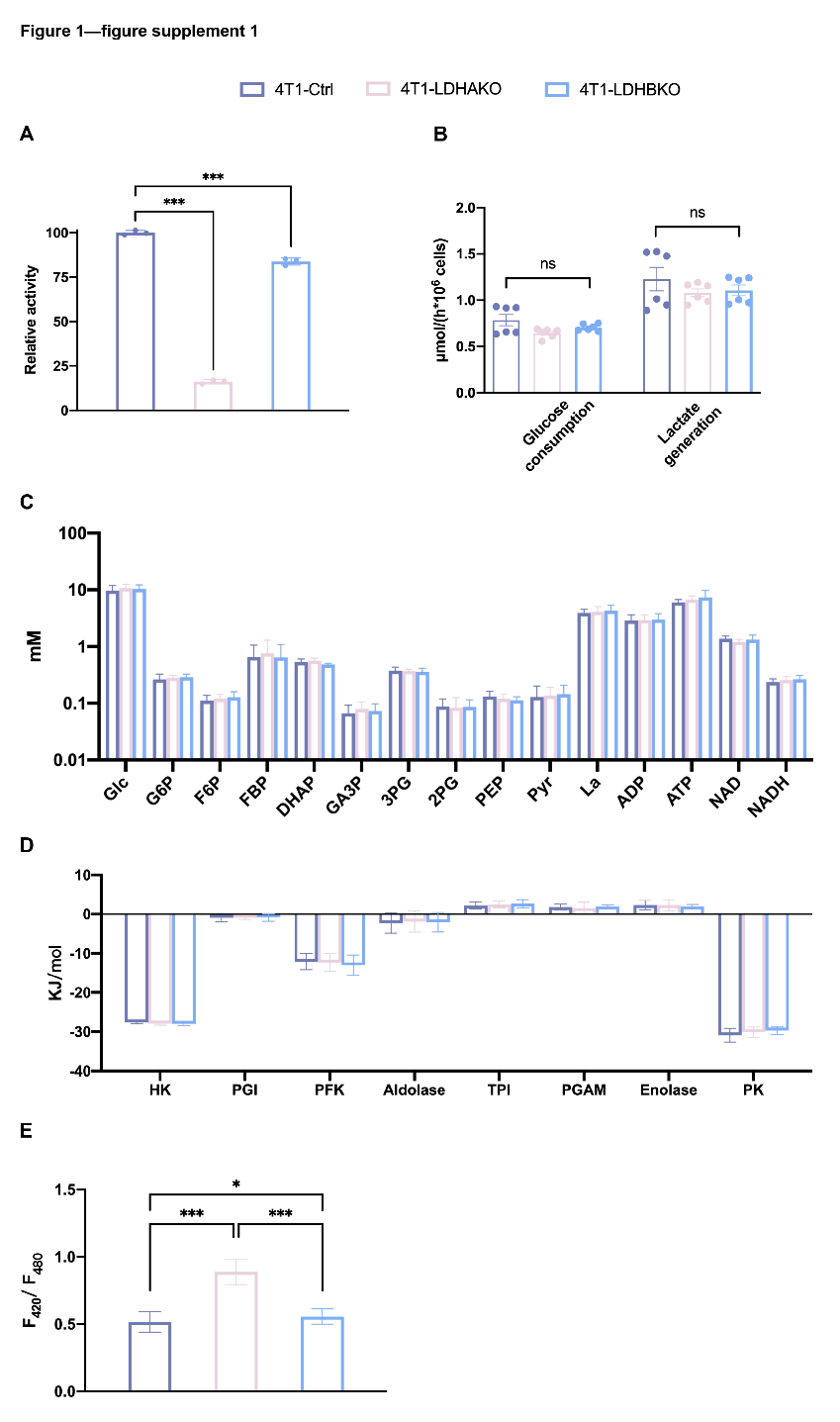

**Figure 1—figure supplement 1. The effects of LDHA or LDHB knockout on LDH activity and on glycolysis in 4T1 cells.** （A）Relative LDH activity in the cell lysate of 4T1-Ctrl, 4T1-LDHAKO, and 4T1-LDHBKO. (B) Glucose consumption rate and lactate generation rate. Cells were cultured in complete RPMI-1640 medium in a CO_2_ incubator for 6 hours, and then the medium concentrations of glucose and lactate were determined as described in Materials and Methods. (C) The concentrations of glucose, lactate, and the glycolytic intermediates in cells (Supplementary table 2). (D) ΔG of the reactions in the glycolytic pathways (Supplementary table 3). (E) Free NADH/NAD^+^ in cells represented by ratiometric of SoNar. Data are mean ± SD, *, *P*<0.05, **, *P*<0.01, ***, *P*<0.001.

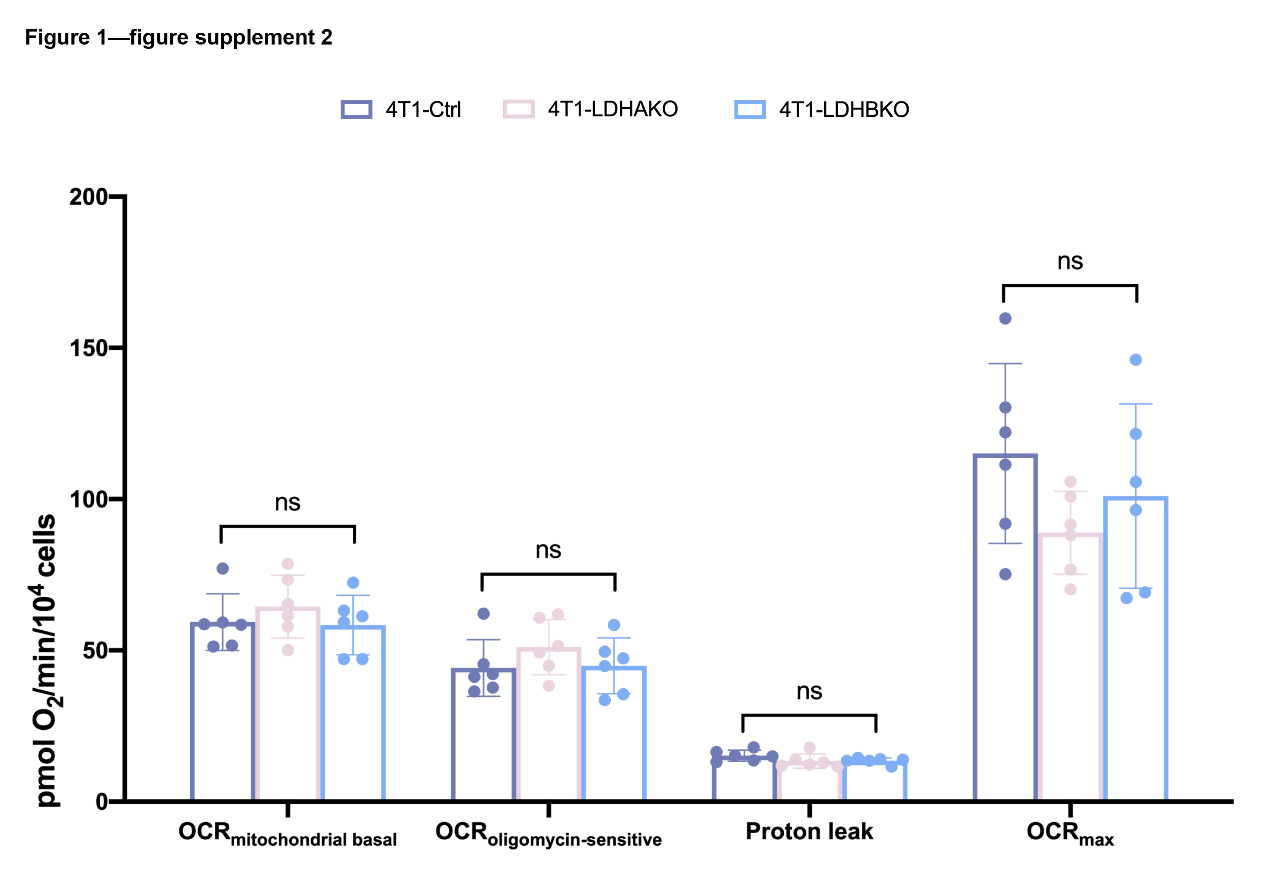
**Figure 1—figure supplement 2. The effect of LDHA or LDHB knockout on OXPHOS in 4T1 cells.** OCR of 4T1-Ctrl, 4T1-LDHAKO, and 4T1-LDHBKO cells. Cells were cultured in complete RPMI-1640 medium in a CO_2_ incubator for 6 hours, and OCR measured as described in Materials and Methods. Data are mean ± SD, *, *P*<0.05, **, *P*<0.01, ***, *P*<0.001.

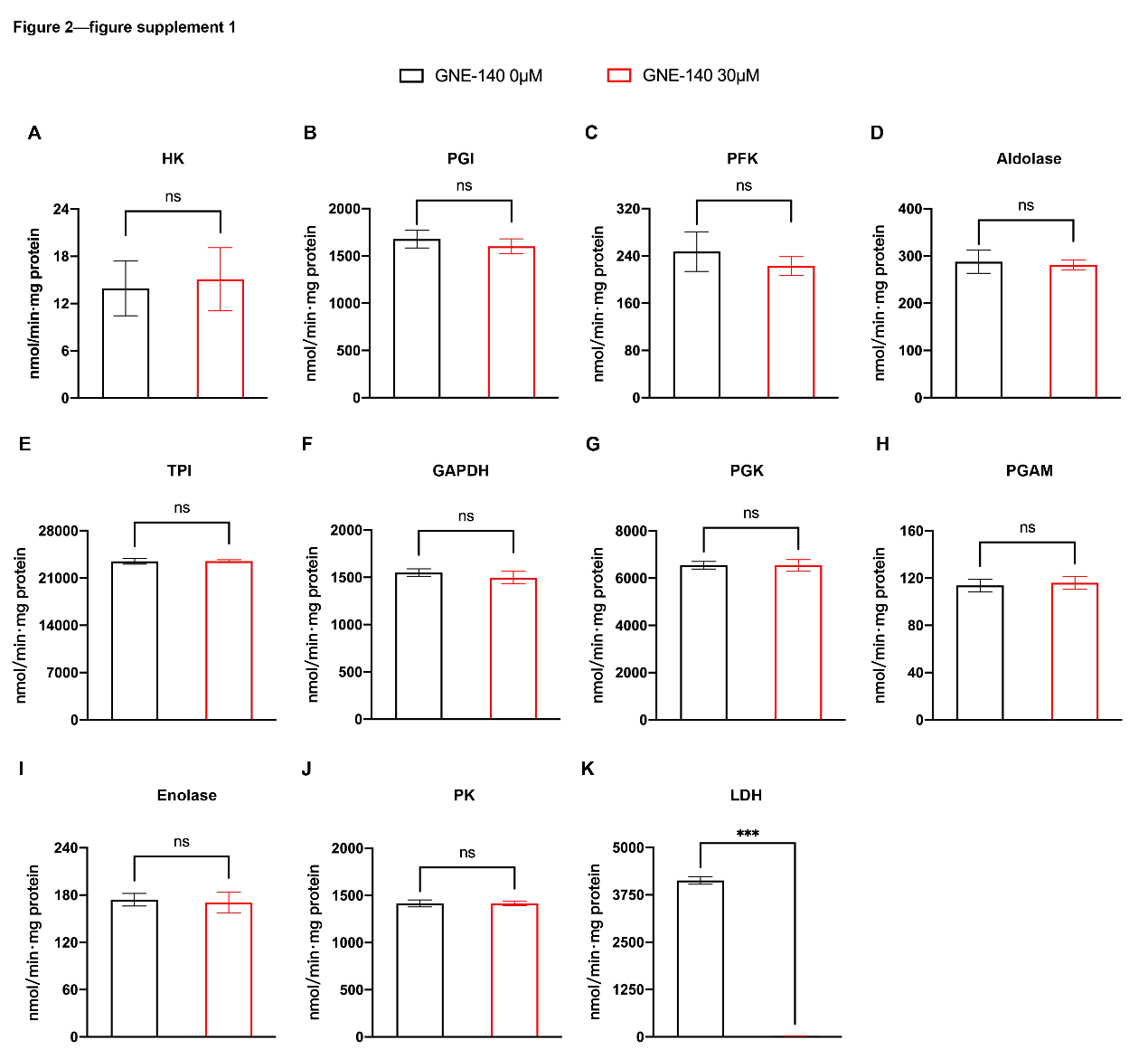

**Figure 2—figure supplement 1. The effect of GNE-140 on glycolytic enzyme activity in HeLa-LDHBKO cells.** (A-K) Glycolytic enzyme activities, including HK, PGI, PFK, aldolase, TPI, GAPDH, PGK, PGAM, Enolase, PK, and LDH. The activity of each glycolytic enzyme was measured with or without the addition of 30 μM GNE-140, as described in Materials and Methods. Data are mean ± SD, *, *P*<0.05, **, *P*<0.01, ***, *P*<0.001.

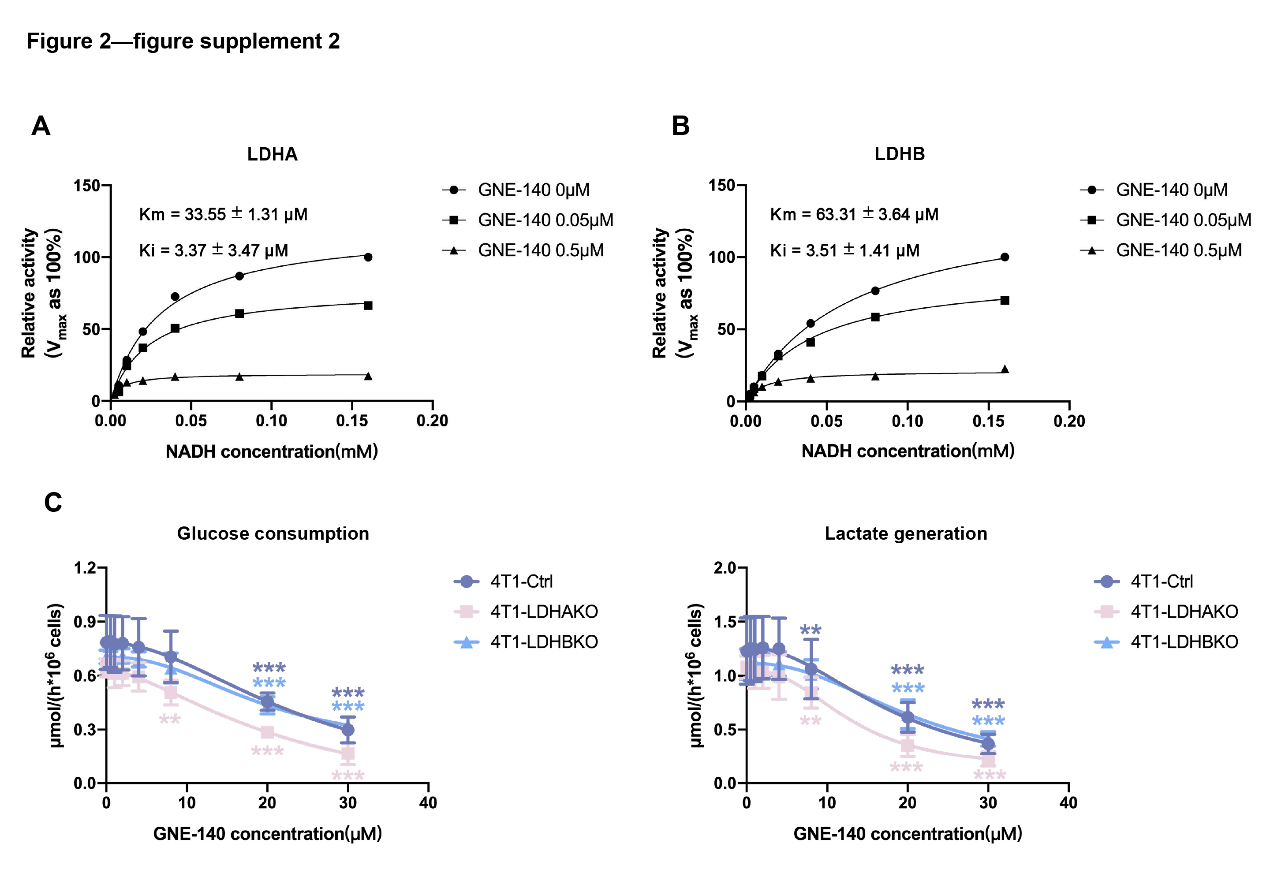

**Figure 2—figure supplement 2. The effect of GNE-140 on LDH activity and on glycolysis in 4T1 cells.** (A & B) Ki of GNE-140 toward LDHA and LDHB. The Ki values were determined as described in Materials and Methods. (C) The effect of GNE-140 on cellular glucose consumption and lactate generation. Cells were cultured in complete RPMI-1640 medium with or without GNE-140 in a CO_2_ incubator for 6 hours and then the medium concentrations of glucose and lactate were determined as described in Materials and Methods. Data are mean ± SD, *, *P*<0.05, **, *P*<0.01, ***, *P*<0.001.

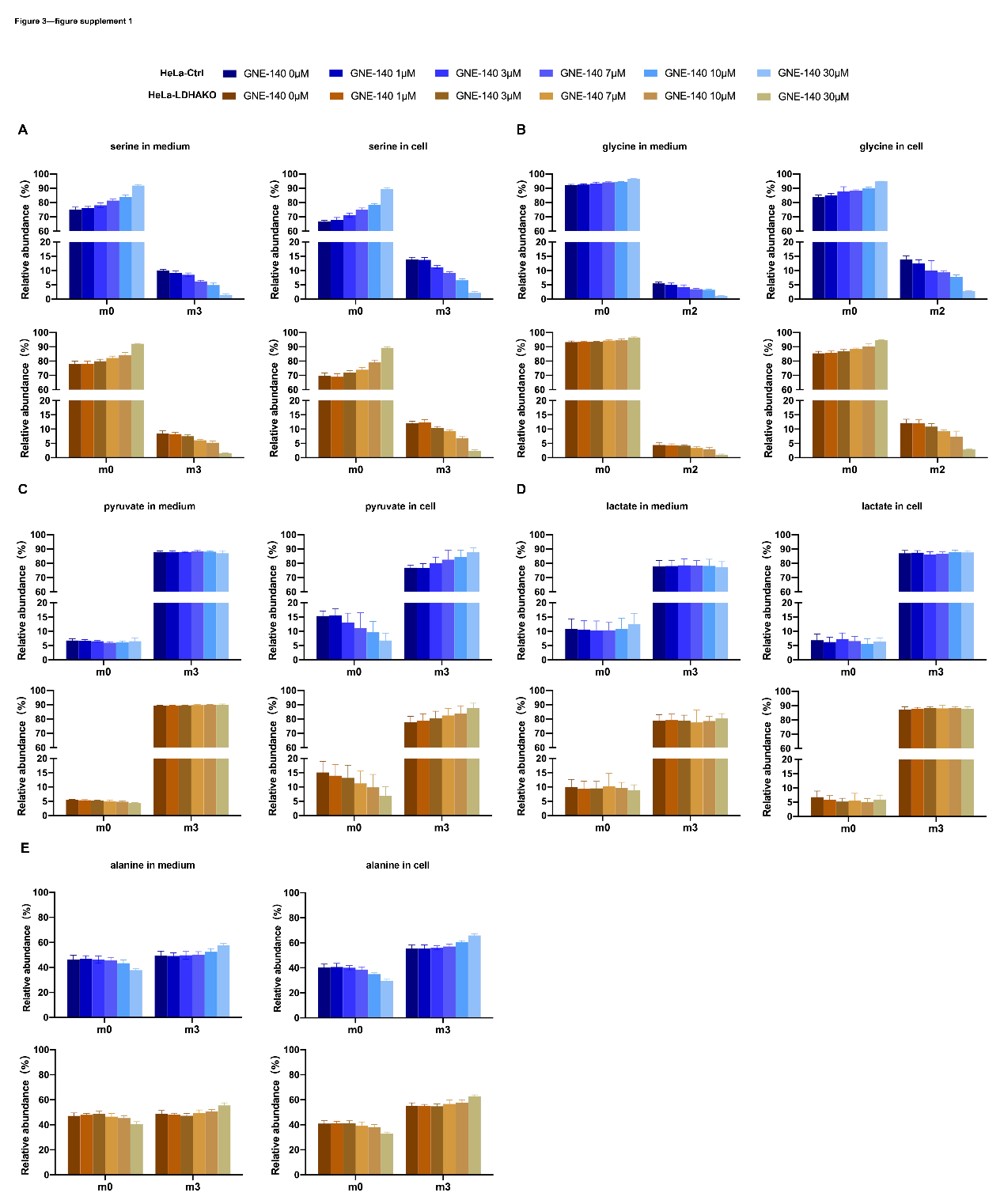

**Figure 3—figure supplement 1. The effect of GNE-140 on the glycolytic pathway in HeLa-Ctrl and HeLa-LDHAKO cells.** Tracing glucose carbon to serine, glycine, pyruvate, lactate, and alanine. Cells were cultured in complete RPMI-1640 medium containing 6 mM [^13^C_6_-glc] with or without GNE-140 in a CO_2_ incubator for 6 hours, and then the percentages of isotopologues in cells and in medium were determined by LC-MS/MS as described in Materials and Methods (Supplementary table 4). (A) Percentages of isotopologues of serine. (B) Percentages of isotopologues of glycine. (C) Percentages of isotopologues of pyruvate. (D) Percentages of isotopologues of lactate. (E) Percentages of isotopologues of alanine. Data are mean ± SD, *, *P*<0.05, **, *P*<0.01, ***, *P*<0.001.

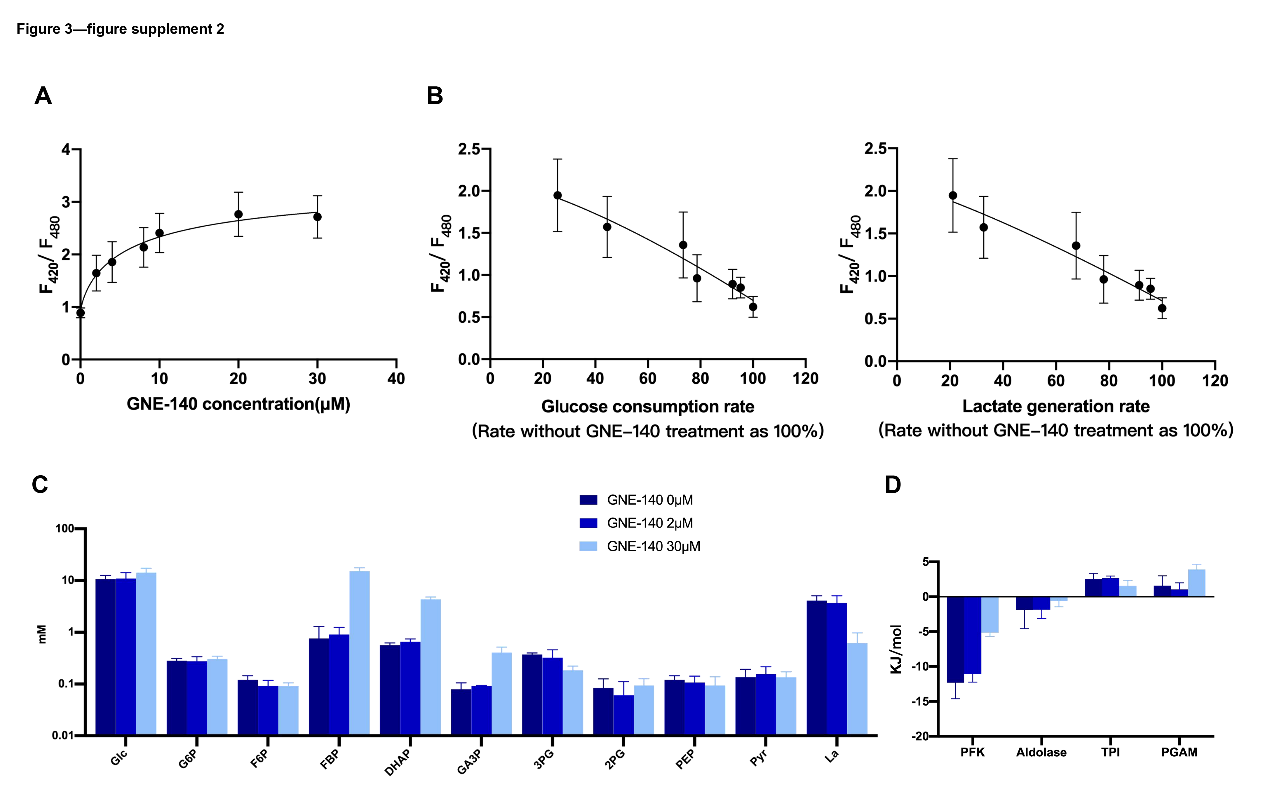

**Figure 3—figure supplement 2. The effect of GNE-140 on the glycolytic pathway in 4T1-LDHAKO cells.** (A) The effect of GNE-140 on the free NADH/NAD^+^. Free NADH/NAD^+^ in cells represented by ratiometric of SoNar. (B) The relationship between the free NADH/NAD^+^ and glycolytic rate. (C) The effect of GNE-140 on the concentrations of glucose, lactate, and the glycolytic intermediates in cells (Supplementary table 5). (D) The effect of GNE-140 on the ΔG of the reactions in the glycolytic pathways in cells (Supplementary table 7). Data are mean ± SD, *, *P*<0.05, **, *P*<0.01, ***, *P*<0.001.

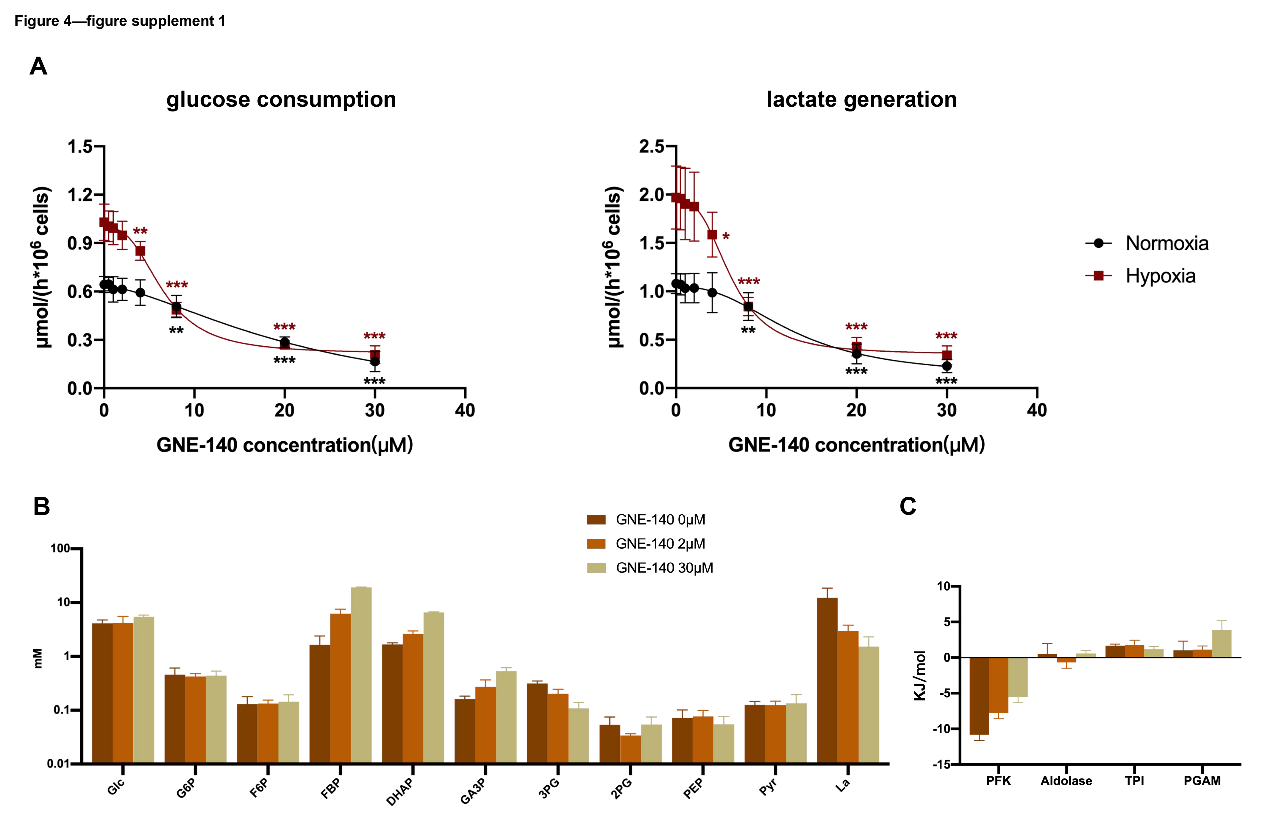

**Figure 4—figure supplement 1. The effect of GNE-140 on the glycolytic pathway in 4T1-LDHAKO cells under hypoxia.** (A) The glucose consumption rate and lactate generation rate in cells under normoxia and hypoxia (1% oxygen) and the response to GNE-140. (B) The effect of GNE-140 on the concentrations of glucose, lactate, and the glycolytic intermediates in cells under hypoxiam (Supplementary table 6). (C) The effect of GNE-140 on the ΔG of the reactions in the glycolytic pathways in cells under hypoxia (Supplementary table 8). Data are mean ± SD, *, *P*<0.05, **, *P*<0.01, ***, *P*<0.001.

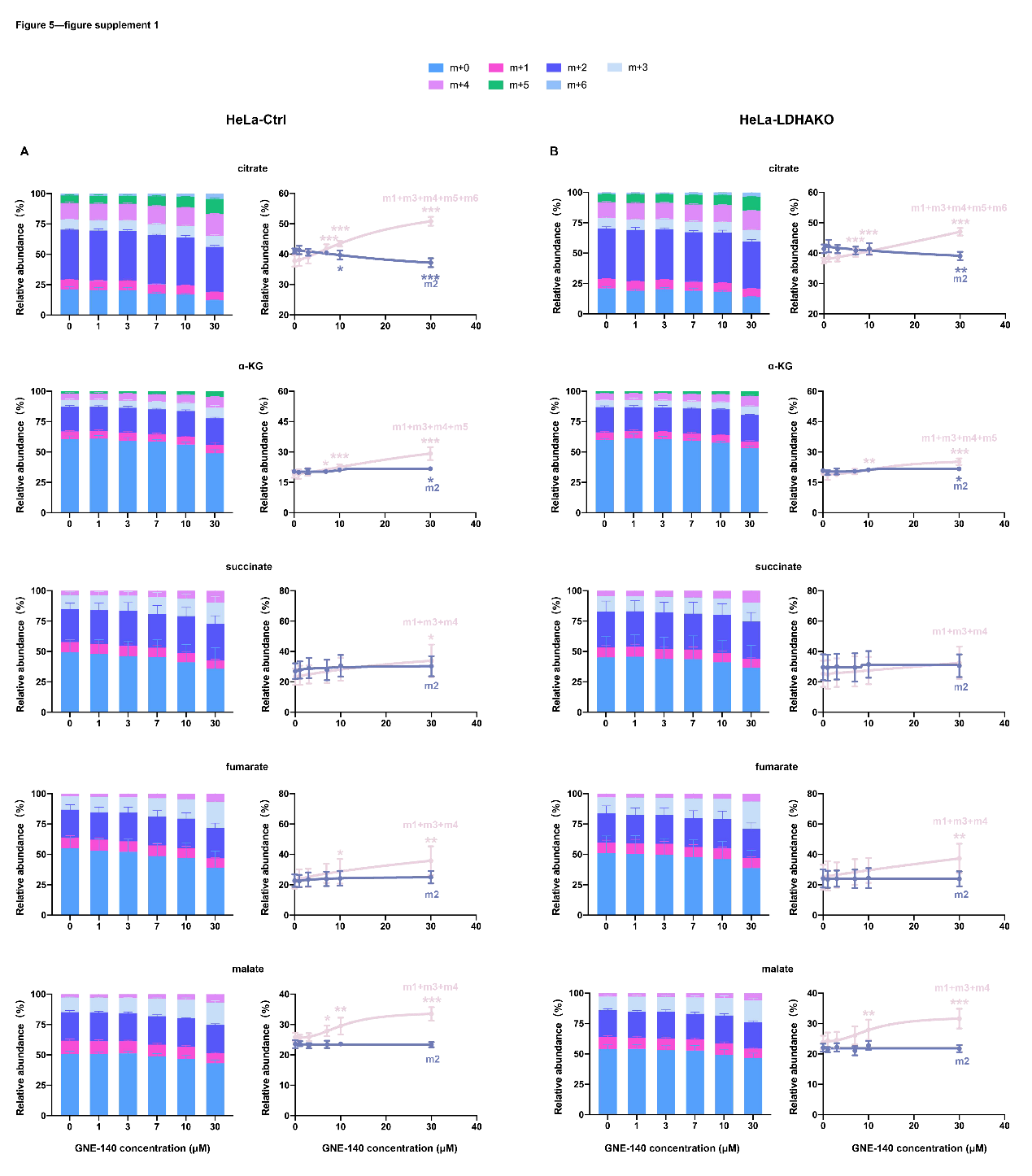

**Figure 5—figure supplement 1. The effect of GNE-140 on glucose carbon into TCA cycle in HeLa-Ctrl and HeLa-LDHAKO cells.** Tracing glucose carbon to TCA cycle intermediates (citrate, α-KG, succinate, fumarate, malate). Cells were cultured in complete RPMI-1640 medium containing 6 mM [^13^C_6_-glc] with or without GNE-140 in a CO_2_ incubator for 6 hours, and then the percentages of isotopologues of the TCA cycle intermediates in cells were determined by LC-MS/MS as described in Materials and Methods. (A) The total ^13^C labeling and the isotope labeling pattern of the TCA cycle intermediates in HeLa-Ctrl cells, including m2 isotopologues% and the sum of other isotopologues% (m1 + m3 + m4 + m5 + m6 for citrate, m1 + m3 + m4 + m5 for α-KG, m1 + m3 + m4 for succinate/fumarate/malate). (B) The total ^13^C labeling and the isotope labeling pattern of the TCA cycle intermediates in HeLa-LDHAKO cells, including m2 isotopologues% and the sum of other isotopologues% (m1 + m3 + m4 + m5 + m6 for citrate, m1 + m3 + m4 + m5 for α-KG, m1 + m3 + m4 for succinate/fumarate/malate). Data are mean ± SD, *, *P*<0.05, **, *P*<0.01, ***, *P*<0.001.

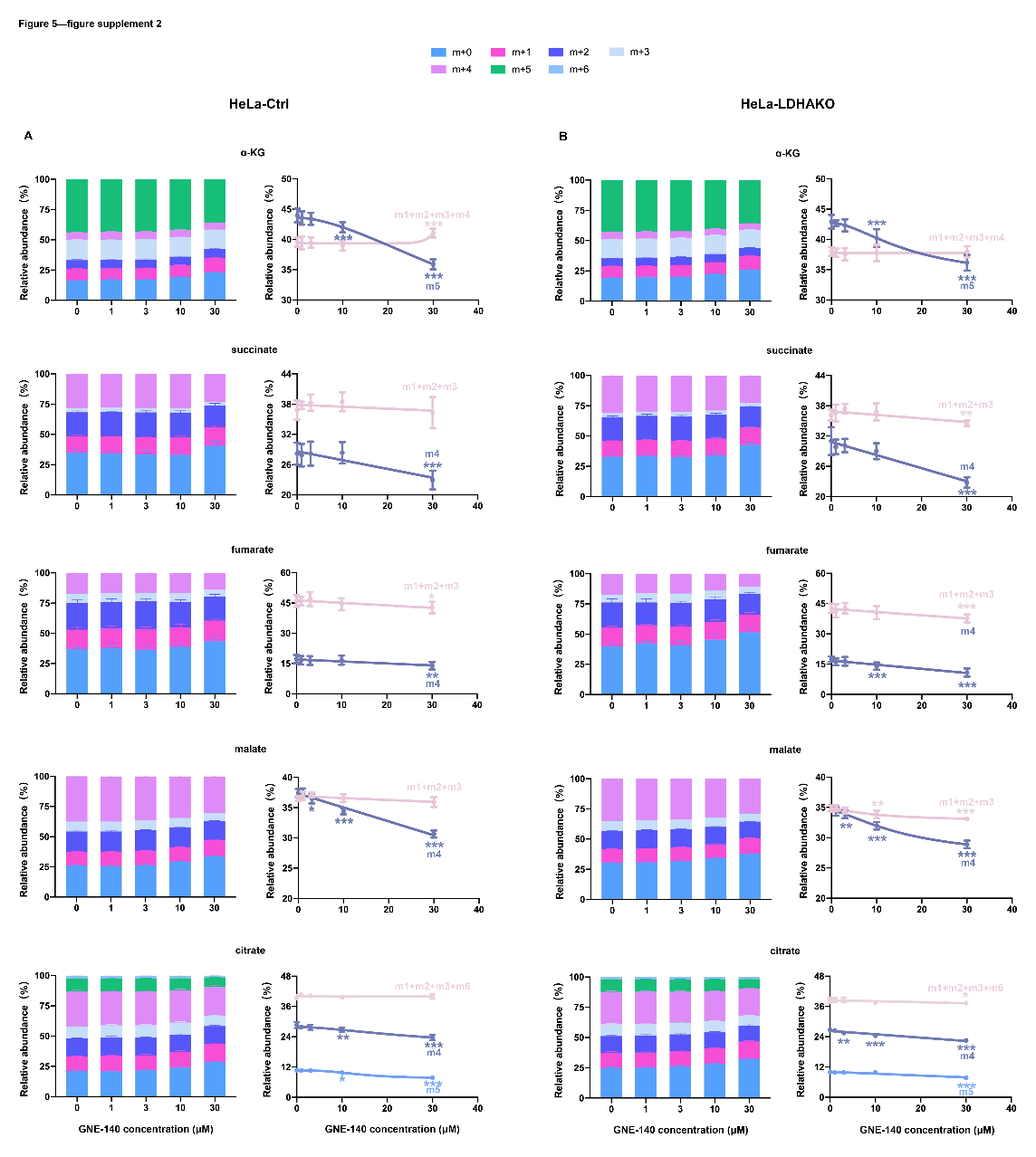

**Figure 5—figure supplement 2. The effect of GNE-140 on glutamine carbon into TCA cycle in HeLa-Ctrl and HeLa-LDHAKO cells.** Tracing glutamine carbon to TCA cycle intermediates (citrate, α-KG, succinate, fumarate, malate). Cells were cultured in complete RPMI-1640 medium containing 2 mM [^13^C_5_-gln] with or without GNE-140 in a CO_2_ incubator for 6 hours, and then the percentages of isotopologues of the TCA cycle intermediates in cells were determined by LC-MS/MS as described in Materials and Methods. (A) The total ^13^C labeling and the isotope labeling pattern of the TCA cycle intermediates in HeLa-Ctrl cells, including m5 α-KG%, m4 succinate%, m4 fumarate%, m4 malate%, m4 citrate%, m5 citrate%, and the sum of other isotopologues% (m1 + m2 + m3 + m4 for α-KG, m1 + m2 + m3 for succinate/fumarate/malate, m1 + m2 + m3 + m6 for citrate). (B) The total ^13^C labeling and the isotope labeling pattern of the TCA cycle intermediates in HeLa-LDHAKO cells, including m5 α-KG%, m4 succinate%, m4 fumarate%, m4 malate%, m4 citrate%, m5 citrate%, and the sum of other isotopologues% (m1 + m2 + m3 + m4 for α-KG, m1 + m2 + m3 for succinate/fumarate/malate, m1 + m2 + m3 + m6 for citrate). Data are mean ± SD, *, *P*<0.05, **, *P*<0.01, ***, *P*<0.001.

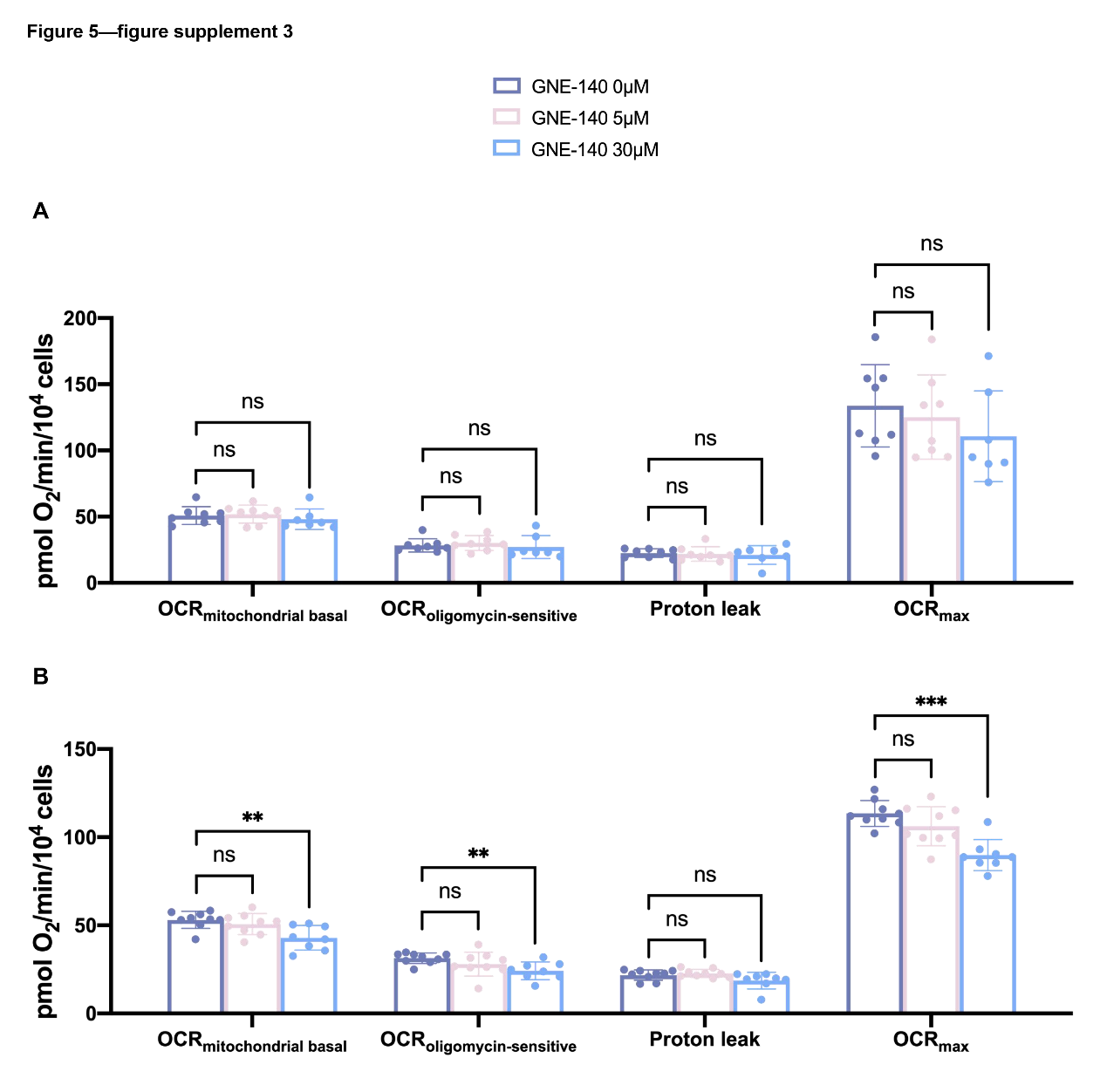

**Figure 5—figure supplement 3. The effect of GNE-140 on OXPHOS in HeLa-Ctrl and HeLa-LDHAKO cells.** Cells were cultured in complete RPMI-1640 medium with or without GNE-140 in a CO_2_ incubator for 6 hours, and OCR measured as described in Materials and Methods. (A) OCR of HeLa-Ctrl cells. (B) OCR of HeLa-LDHAKO cells. Data are mean ± SD, *, *P*<0.05, **, *P*<0.01, ***, *P*<0.001.

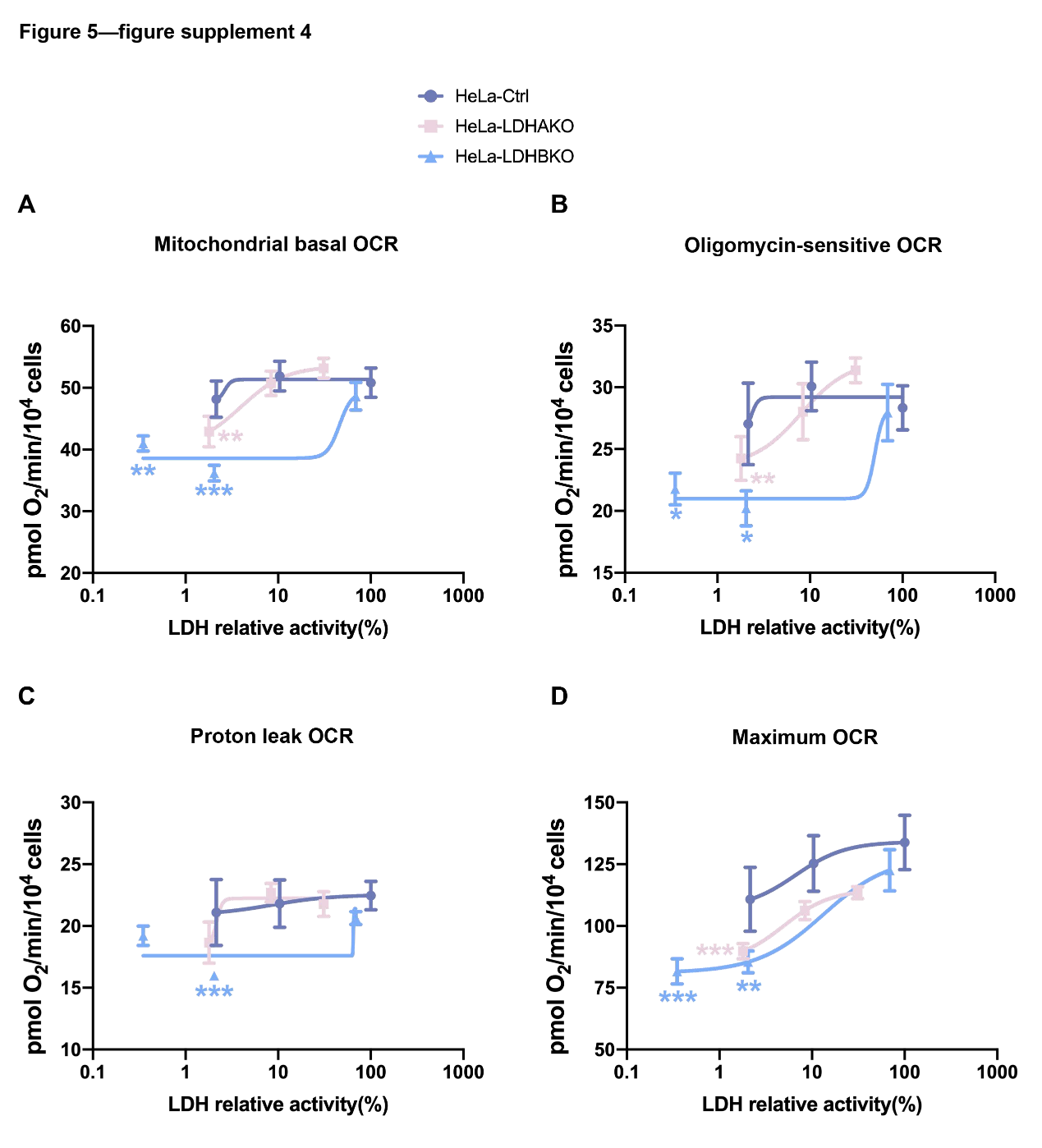

**Figure 5—figure supplement 4. The relationship between the OXPHOS and LDH relative activity in HeLa cells.** Calculate the remaining LDH relative activity of HeLa-Ctrl, HeLa-LDHAKO, and HeLa-LDHBKO cells with different concentrations of GNE-140, and draw the corresponding OCR-LDH relative activity curve. (A) Mitochondrial basal OCR-LDH relative activity curve. (B) Oligomycin-sensitive OCR-LDH relative activity curve. (C) Proton leak OCR-LDH relative activity curve. (D) Maximum OCR-LDH relative activity curve. Data are mean ± SD, *, *P*<0.05, **, *P*<0.01, ***, *P*<0.001.

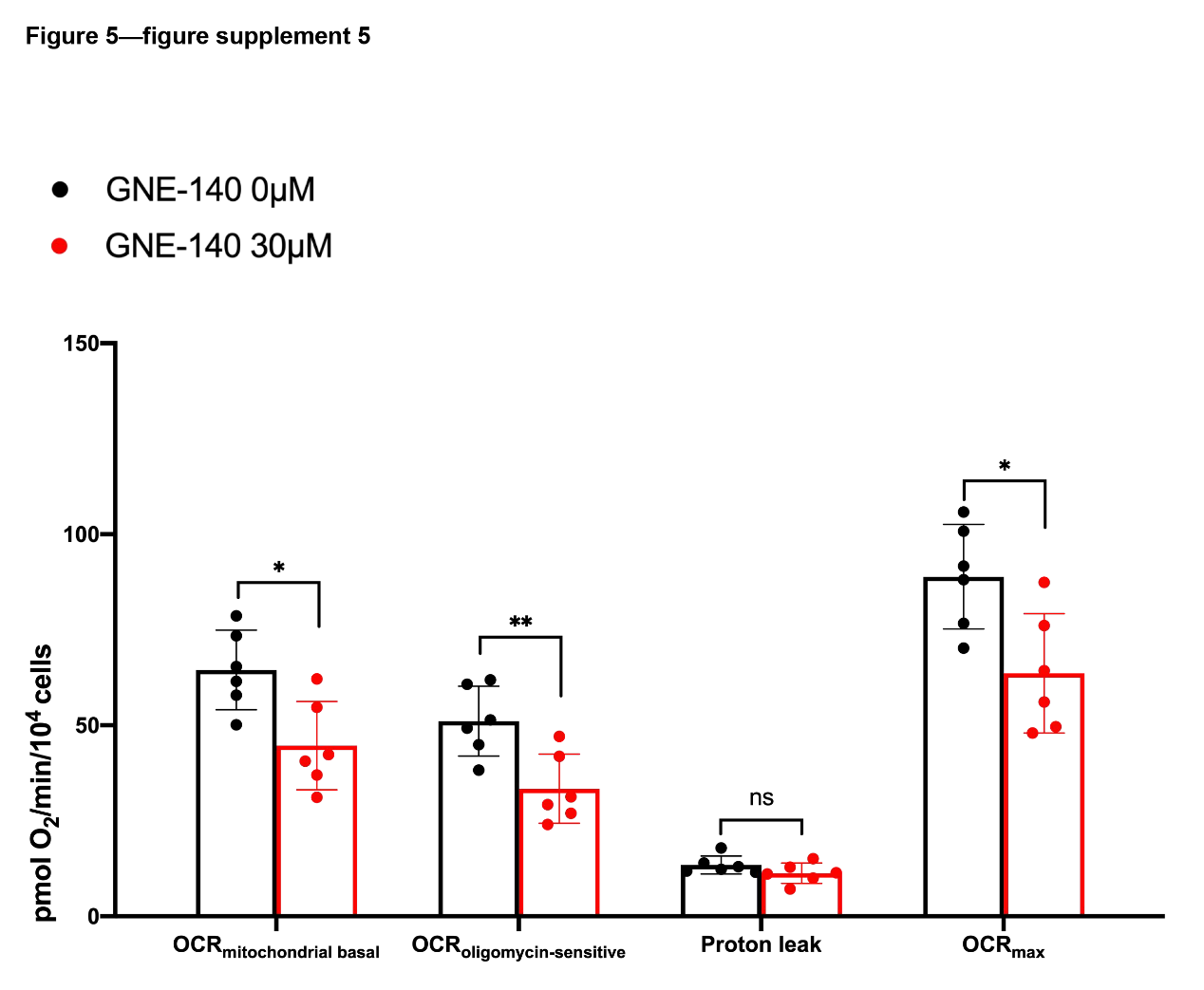

**Figure 5—figure supplement 5. The effect of GNE-140 on OXPHOS in 4T1-LDHAKO cells.** OCR of 4T1-LDHAKO cells. Cells were cultured in complete RPMI-1640 medium with or without GNE-140 in a CO_2_ incubator for 6 hours, and OCR measured as described in Materials and Methods. Data are mean ± SD, *, *P*<0.05, **, *P*<0.01, ***, *P*<0.001.

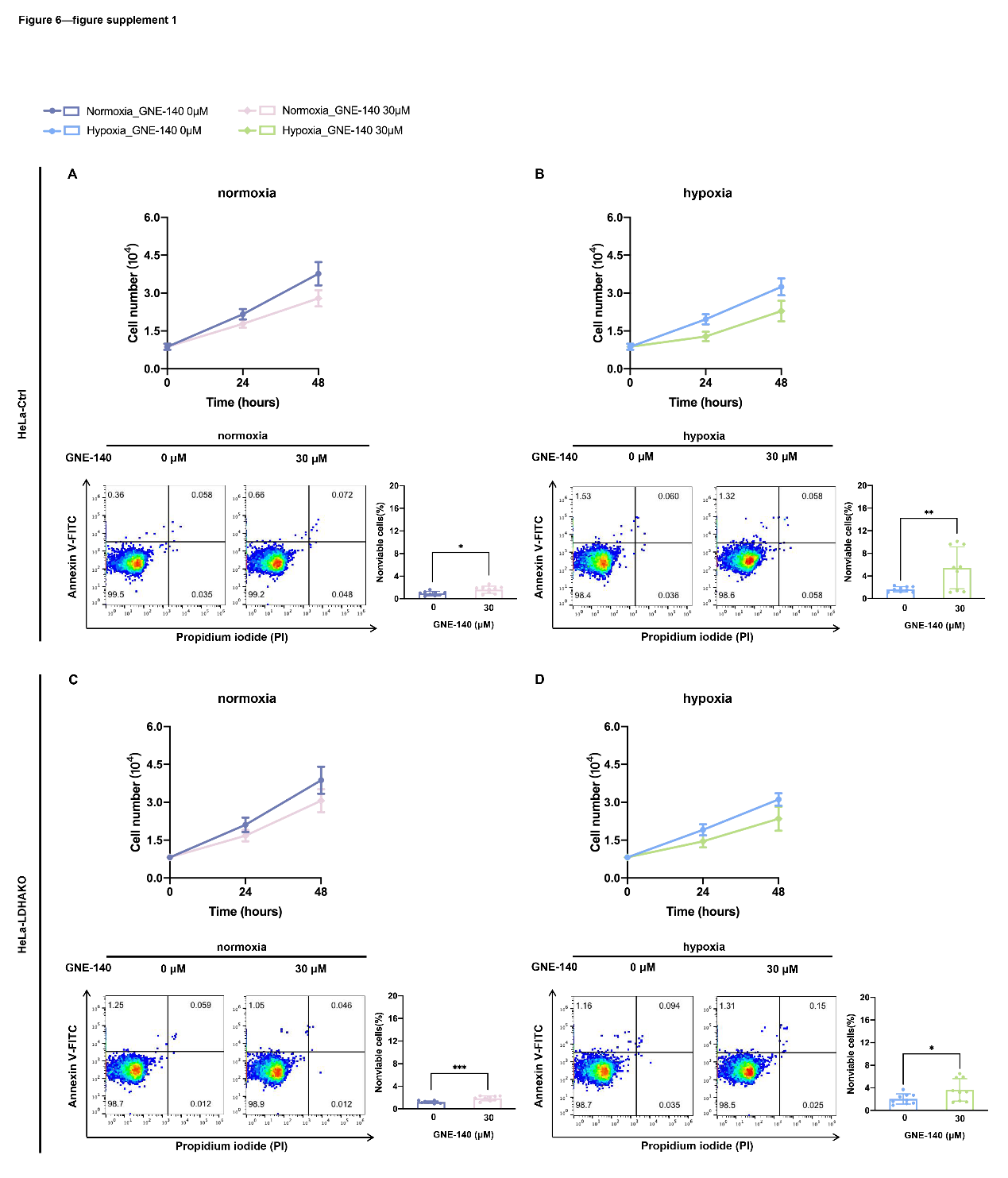

**Figure 6—figure supplement 1. The effect of GNE-140 on cell survival in HeLa-Ctrl and HeLa-LDHAKO cells under normoxia and hypoxia.** (A & B) Cell growth curves and cell death assays under normoxia and hypoxia of HeLa-Ctrl cells. (C & D) Cell growth curves and cell death assays under normoxia and hypoxia of HeLa-LDHAKO cells. Data are mean ± SD, *, *P*<0.05, **, *P*<0.01, ***, *P*<0.001.
